# GDEE: A Structure-Based Platform for Gene Discovery and Enzyme Engineering

**DOI:** 10.1101/2025.09.09.675117

**Authors:** Caio S. Souza, João P.G. Correia, Isabel Rocha, Diana Lousa, Cláudio M. Soares

**Affiliations:** Instituto de Tecnologia Química e Biológica António Xavier, Universidade Nova de Lisboa, Av. da República, 2780-157 Oeiras, Portugal

## Abstract

Enzymes are extensively utilized for catalyzing the production of value-added compounds in diverse sectors with significant economic impact. However, challenges such as poor protein expression, low catalytic activity, substrate/co-factor limitations, and toxicity of final products limit the efficiency of these biocatalysts. Protein engineering offers a solution by re-designing enzyme catalytic properties to enhance biosynthetic pathways. In this study, we present an automated platform for gene discovery and enzyme engineering, aimed at overcoming these bottlenecks. The GDEE platform finds and optimizes enzymes responsible for rate-limiting steps in biosynthetic pathways, with objectives ranging from improving catalytic efficiency to enabling novel transformations. The platform operates through four key steps: an initial sequence step that either sources natural enzyme sequences or generates mutant variants, followed by an atomistic protein structure prediction step, a docking step, and a binding energy evaluator, obtaining a set of variant enzyme sequences that are prioritized for experimental validation. Additionally, machine learning-based re-scoring of binding poses to improve binding affinities significantly enhances the accuracy of predicting the effect of mutations on binding, highlighting its potential in metabolic engineering applications.

## Introduction

Poor enzyme expression, low catalytic levels, low cofactor or substrate concentrations, toxicity toward the end product, and other variables can all contribute to low performance of natural or novel pathways in microorganisms (Du et al., 2011; Geraldi et al., 2021). Improving enzymatic DNA transcription has been the most popular technique for overcoming some of these challenges. However, other difficulties such as cofactor availability, low substrate affinity, or inhibition of catalysis, cannot be addressed by increasing enzyme concentration. Enzyme engineering to optimize metabolic pathways is an effective strategy for re-designing the catalytic characteristics to favor a certain flow (Pleiss, 2011). It is feasible to enhance the catalytic properties by increasing substrate affinity and selectivity, or to eliminate inhibition (L. Wang et al., 2022). Several techniques and protocols have been created, in order to do enzyme engineering, employing random mutagenesis and selection via direct evolution, or hybrid methods where rational design is employed to reduce the reliance on widespread random mutagenesis (Bloom et al., 2005).

Computational tools are also important in enzyme engineering. When the structure of the enzyme is understood, rational design approaches may be used. These computational techniques use enzyme structural knowledge to predict the impact of modifications of individual amino acid residues on enzyme activity. In rational protein engineering, several modeling methods can be utilized, ranging from highly detailed low throughput methods to methods that allow high-throughput analysis by adding approximations. In the first group, we have Quantum Mechanics (QM) calculations and combined Quantum Mechanics and Molecular Mechanics (QM/MM) techniques. These approaches are particularly meticulous since they treat the enzyme active site at the quantum level and allow for an analysis of reaction mechanisms as well as prediction of activation barriers. These approaches, however, are not sufficiently fast to be used in high-throughput calculations (Kiss et al., 2013; Marcheschi et al., 2012; Ranaghan & Mulholland, 2010).

Molecular mechanics (MM) based free energy calculations and other MM-based methods, such as meta dynamics, can also be used to calculate free energy differences and thus predict how mutations affect the binding of a given substrate or reaction intermediate, but these methods are also computationally demanding (Ranaghan & Mulholland, 2010). Molecular docking approaches are much faster due to the numerous approximations introduced (such as assuming the protein rigid or semi-rigid and not employing explicit solvent) and predict the binding free energy of a ligand to a protein by applying empirical free energy functions (Morris & Lim-Wilby, 2008). Docking techniques have been used successfully in rational enzyme design (Sirin et al., 2014), and, although not as thorough as QM-based approaches, these methods are faster and may be employed in a high-throughput manner. AutoDock Vina is especially well-suited for this task, in view of its calculation efficiency, allowing it to run faster without sacrificing prediction of binding poses and energies (Trott & Olson, 2010). To apply docking approaches in a high-throughput manner, one must have an automated way of generating and preparing mutant enzyme structures as well as executing docking calculations.

To overcome this problem, we developed an automated platform that combines several structural bioinformatic tools that can be used to perform structure-based (i) enzyme engineering by mutating specific residues of a given active site to optimize the enzyme transformation, or (ii) gene discovery, where natural enzymes are scanned to find those that are more prone to catalyze the target reaction. This platform will predict which of the candidate enzymes has a better affinity for the molecule of interest (substrate, intermediate or product) given an initial set of enzymes (which can be *in silico* generated mutants of the same enzyme or natural occurring variants). Rather than calculating catalytic efficiency directly, which is computationally demanding and often unfeasible for large variant libraries due to the complexity of modeling full enzymatic reaction mechanisms, this platform uses predicted binding affinity as a practical proxy. Binding affinity indicates how strongly an enzyme can bind to its target molecule, which is often a crucial step for the enzyme to perform its functions, and can be efficiently estimated for many candidates. This approach enables effective screening in medium-to high-throughput workflows (Chu & Wang, 2016; Su et al., 2021; Xia et al., 2021).

### Overview

The gene discovery and enzyme engineering pipeline consist of a sequence of steps aiming at designing putatively more efficient variants of enzymes. The Pipeline steps are grouped into four main steps: an initial sequence step that either sources natural enzyme sequences or generates mutant variants, followed by an atomistic protein structure prediction step, a docking step, and a binding energy evaluator. The platform, at this stage, assumes that a protein with a set of mutations that increases the enzyme affinity to substrates, products or reaction intermediates is prone to have better kinetic parameters. Thus, with this pipeline, it is possible to generate a set of variant enzymes and filter which sequences are putative candidates for experimental expression and enzymatic testing.

Before starting to engineer an enzyme, it is crucial to have a comprehensive understanding of the structural and dynamic foundations of enzymatic function. The availability of an experimental structure remains highly valuable for successful engineering, as it provides precise insights into key regions such as the binding pocket, where mutations can significantly influence the binding affinity of the target ligands. In cases where an experimental structure is not available, computational protein structure prediction can be used to generate reliable models as templates. Notably, with recent advances like AlphaFold3 (Abramson et al., 2024), it is now possible to predict enzyme structures in complex with their ligands, offering enhanced insights into enzyme-substrate interactions. This capability greatly improves confidence in structure-based engineering efforts, although the accuracy and quality of these predicted enzyme-ligand models still play a critical role in determining the success of subsequent engineering.

### GDEE platform

The GDEE platform is developed as a Python package composed of 8 distinct modules, as illustrated in Figure 1. Each module serves a specific role within the enzyme engineering or gene discovery pipeline. The Pipeline module orchestrates the pipeline workflow and data persistence. The Variant module gathers protein sequence variants using different strategies, including natural or mutant sequences. The modeling module predicts the three-dimensional structure of each variant and assesses the quality of the modeled structures. The Evaluator module performs protein-ligand docking and calculates binding affinities. The Measurement module computes distances between selected atoms of the protein and ligand to support filtering of docking poses. The analysis module filters docking poses based on user defined metrics and ranks the enzyme variants accordingly. The Platform module manages execution modes, supporting both single-threaded and distributed workflows for scalability. Finally, the database module handles all data persistence, utilizing SQLite3 to store and organize data for effective post-processing and re-analysis.

**Figure 1.**
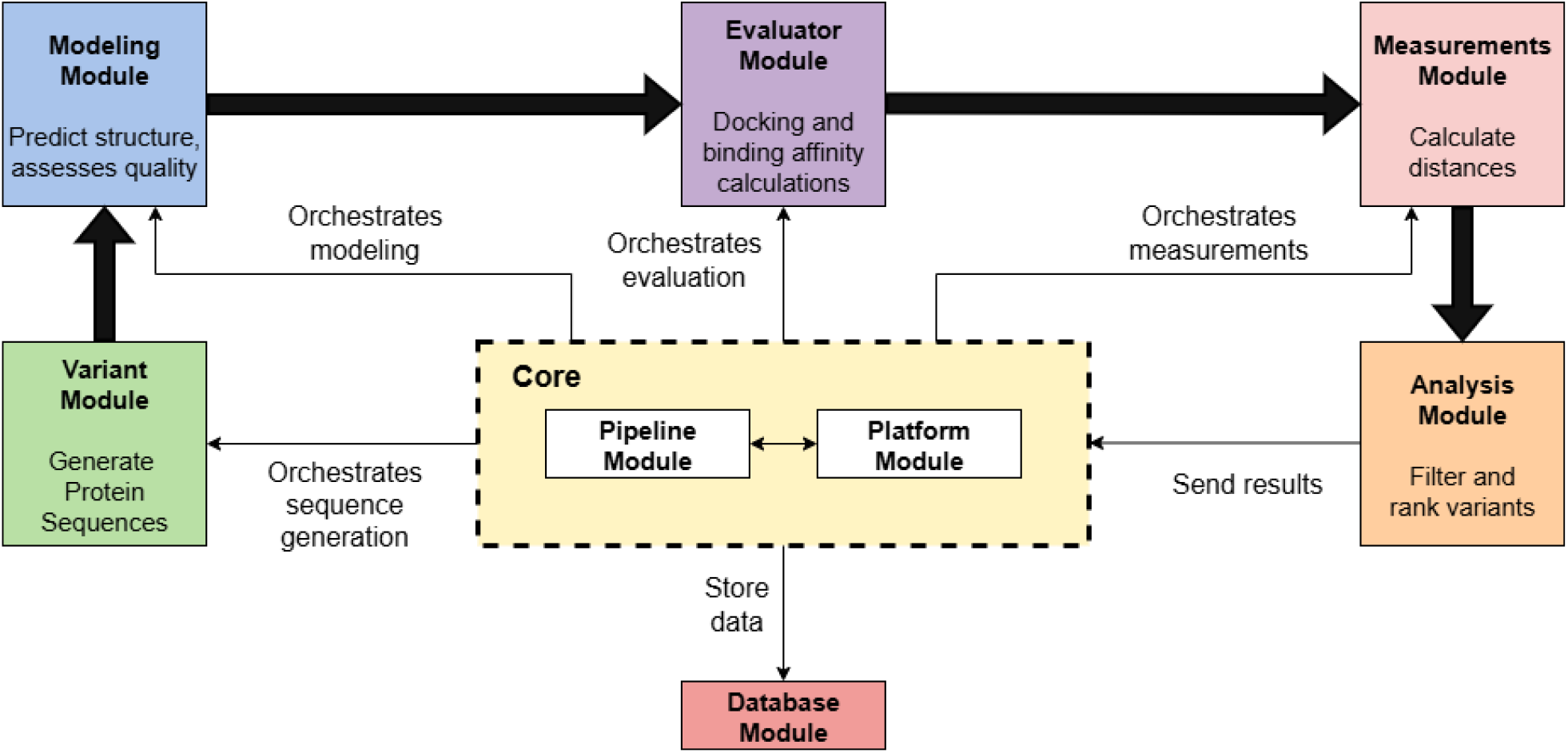
High-level diagram illustrating the architecture and data flow of the platform. The Core functions as the central orchestrator coordinating and scheduling task execution across specialized modules in a linear pipeline workflow – Variant, Modeling, Evaluator, Measurement, and Analysis – each dedicated to specific tasks on protein variants. The Variant module generates amino acid sequence variants using different strategies; the Modeling module predicts 3D structures and assesses their quality; the Evaluator module performs docking and binding affinity calculations; the Measurement module calculates distances between selected protein and ligand atoms to assist in filtering; and the Analysis module applies filters and ranks variants accordingly. Data flows between processing modules are represented by bold arrows, while narrow arrows indicate the Core orchestration and management of operations. All intermediate and final results are centralized in the Database module, providing efficient data storage and retrieval for downstream analysis.

The package is designed with an object-oriented and modular architecture that facilitates extensibility. New algorithms and functionalities can be seamlessly integrated, allowing customization and growth of the platform.

External software dependencies play a key role in the pipeline. The modeling module uses Modeller (Šali & Blundell, 1993) via its Python package integration for structure prediction. Docking and affinity calculations rely on AutoDock Vina (Trott & Olson, 2010) or Smina (Koes et al., 2013). Parallel processing is implemented using MPI4Py (Dalcin & Fang, 2021), a Python interface to the Message Passing Interface (MPI), enabling efficient data parallelism across computational resources.

All results and intermediate data are stored in a SQLite3 database, ensuring organized data management and facilitating downstream analysis and reproducibility.

### Gene Discovery and Enzyme Engineering pipeline

The initial input for each job is the structure of the template enzyme and the structure of one or more ligand molecules. The molecular structures of the protein and ligand are expected to be in a PDB (Protein Data Bank) and PDBQT file format, respectively. After providing the initial data it is possible to create a new job that will proceed with the pipeline steps. The pipeline starts with the generation of a set of candidate enzyme sequences to be evaluated. Multiple homology models are built and optimized for each sequence variant using the user-provided template structure. The final models undergo molecular docking calculations with a ligand that represents the reaction’s limiting step (usually the reaction intermediate or substrate). Finally, the sequences are ranked according to the binding affinity computed in the docking step.

#### Sequence generation step

The gene discovery and the enzyme engineering features differ only in the sequence generation step. For enzyme engineering, sequences are automatically created by sampling the mutation space of user-selected residues or by generating all possible combinations of amino acids for each residue position selected. In the case of sampling, a user-defined number of amino acid residue substitutions is chosen based on the scoring of a substitution matrix for each individual annotated site. Mutations may be conservative (lowest substitution cost), non-conservative (highest substitution cost) or completely random (substitution matrix is ignored). The platform can generate new sequences by applying groups of mutations of varying sizes, allowing simultaneous substitutions of one, two, three, or more amino acids according to the desired mutations count. For gene discovery, variant sequences are submitted to the platform as a FASTA file. These variant sequences are pre-selected based on their similarity to the template enzyme, identified through BLAST searches against sequence databases. The mutation space scales exponentially with the number of residues to mutate, and, for this reason, the GDEE platform was created to allow automated and high-throughput generation and testing of thousands of protein sequences.

#### Model generation and optimization

Homology models are built and optimized by Modeller with the user-provided enzyme structure serving as a template. Models are built using the methods present in the program. Special restraints may be applied in the modelling phase, such as an optimization radius and an optimization level. The optimization radius defines the subset of amino acid residues selected for geometry optimization. This subset can include only the mutated residues (with an optimization radius of 0) or both mutated and neighboring residues. The optimization radius, measured in angstroms, sets a cutoff distance from the mutated residues to select neighboring atoms for optimization. This optimization applies only to side chains, while keeping the backbone fixed to preserve the protein’s overall folding. The user may also choose optimization and refinement levels to the optimization process. There are three optimization levels (0, 1 and 2) implemented. Level 0 refers to the fast refinement available in Modeller with 100 maximum calls of the objective function, level 1 refers to the slow refinement with 200 maximum iteration and level 3 to the Modeller’s very slow refinement with 300 maximum calls of the objective function. The platform will create three times the number of models defined by the user for each of the sequences generated in the previous step, allowing to explore structural diversity, and enabling quality assessment and selection of the best models. To prevent inferior quality models from being passed on to the docking step and then delivered to the user, two structural quality assessment methods, namely VoroMQA (Olechnovič & Venclovas, 2017) and Modeller’s normalized DOPE (Shen & Sali, 2006)) are available in the platform, decreasing the amount of manual curation of the resulting models and improving the performance of the platform by avoiding unnecessary docking calculations. All the models generated are individually scored by each of the quality assessment methods and classified into accepted or rejected. Then, only the best models (based on these scores), according to a user defined number of desired models, are passed on to the docking step. While VoroMQA and the normalized DOPE methods are able to identify significant structural problems, their sensitivity falls short when it comes to detecting subtle structural problems. Consequently, some manual curation remains imperative for refining the resulting models, especially in addressing minor structural anomalies.

#### Docking and binding affinity calculations

Docking and binding affinities calculations between the ligand and the protein are performed on all of the generated variant models that passed through the acceptance criteria by the programs AutoDock Vina or Smina. Binding energies are obtained by docking of each provided ligand structure. The end-user may choose between Vina or Vinardo scoring functions. The initial ligand position is obtained from the user specified PDBQT file without modifications. The exhaustiveness parameter can be defined by the user and random seeds are randomly generated on each run. The box size and center must be provided by the user. The reference binding affinity (binding affinity of the reference protein (WT)) is also obtained by the aforementioned methods. The protein is considered rigid throughout the binding affinity calculations. Protein structures are automatically processed by AutoDockTools (Morris et al., 2009) scripts.

#### Data parallelism

We note that the steps described above can be performed on distributed systems using MPI.As a parallelizable solution, the GDEE platform enables automated high-throughput calculations, significantly enhancing computational efficiency.

#### Storage of data

All data obtained on each step is stored and organized as entries in specific tables in a SQLite database. Data is thus easily accessible to debug the pipeline or to perform extra analysis needed by the end-user, such as filter poses to obtain only those in catalytically relevant orientations.

The database is composed of 7 tables (Evaluations, Measurements, Metrics, Models, Poses, Proteins and Variants). The diagram for the relational model of the database is shown in Figure 2. The table “Proteins” contains the records for each reference protein used. Here we can have information on the name of the protein and the Uniprot ID. The table “Variants” register information on the name of the variant (in enzyme engineering cases, this is the mutation, e.g. A:V220L, and in the gene discovery cases, this contains the Uniprot ID), its sequence, the directory name where the resulting files are stored and a Boolean to indicate whether it is the wild type (or the reference protein in gene discovery cases). It also has the possibility to add a PDB file and code for each variant if the user desires to. The “Evaluations” table links the variants, models, and poses to an evaluation. In addition to these links, there is also information on the ligand name and file used in each evaluation process, the method utilized for the evaluation step and the resulting PDB file. The “Models” table retains information regarding the variant which the model corresponds to, the software used to create the model, the scores for each model, the resulting PDB file and a Boolean to indicate if the model was rejected or not. The information about the docking results is stored in the “Poses” table, which contains the pose index and the associate energy for each evaluation. The two remaining tables “Metrics” and “Measurements” hold information on the distances that are going to be used for the filtering step. The first one, associate the metric’s name to the metric itself and the other links each pose to the metric and its value.

**Figure 2.**
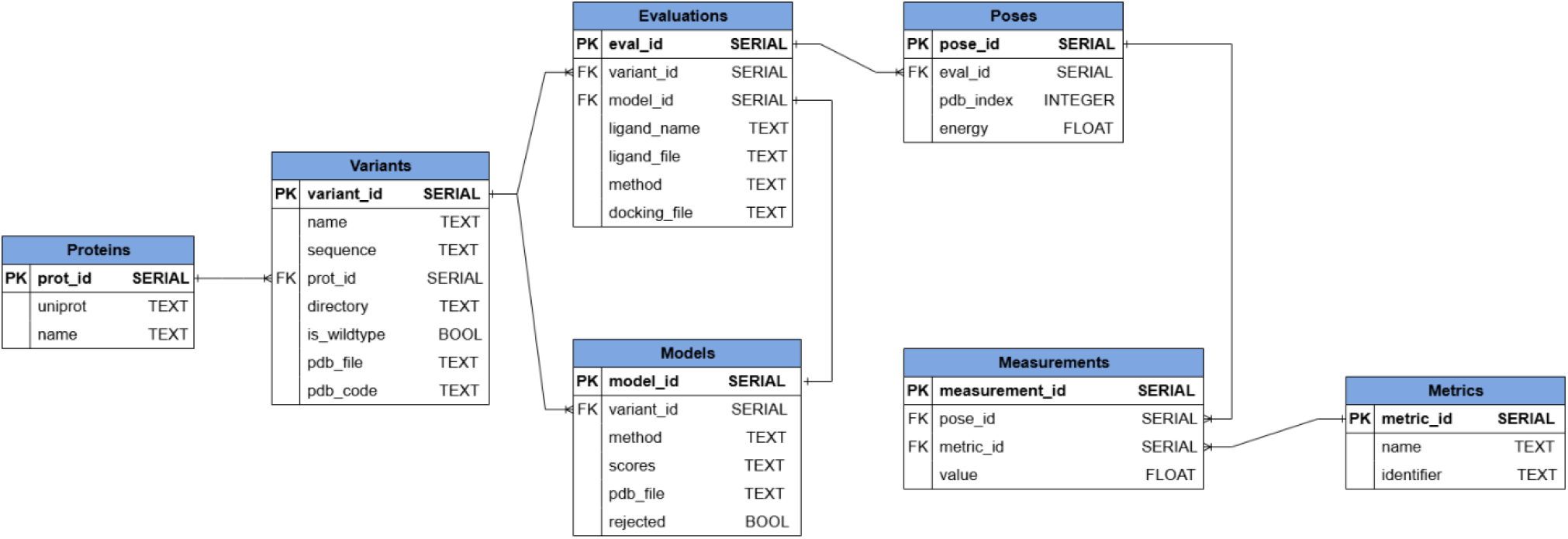
Relational model (entity-relationship diagram) of the platform’s database. This diagram depicts the logical structure and organization of the database used here, including all entities, their principal attributes, and the relationships among them. Entities are represented as tables containing attributes, with primary keys indicated and foreign key relationships explicitly shown. Relationships between entities—such as one-to-many or many-to-many associations—are indicated using standard ERD notation for cardinality. This model captures how resulting data is integrated, supporting robust data storage, retrieval, and linkage across all steps. See text above, for the explanation.

#### Filtering process and ranking step

To ensure that ligands are in a catalytically relevant orientation, the docking results are filtered using distance constraints between atoms from the ligand and the candidate enzyme. The distances are stored in the database mentioned above, so ligand poses relevant to the reaction are easily queried. For cases where more than one pose is accepted in the filtering criteria, the selected ligand pose for each variant sequence is the one with the best binding affinity value.

### Preparation protocols: Gene Discovery

The standard preparation protocol for gene discovery begins with selecting a template structure that provides an appropriate protein fold for the *in-silico* search for enzymes capable of catalysing the target reaction. Notably, the template does not need to catalyse the target reaction itself; it simply needs to offer a compatible structural framework. Once a suitable template is identified, the ability of the active site to accommodate the reaction substrate or intermediate is typically assessed by generating a model of the enzyme–ligand complex. This can be accomplished either through *ab initio* methods such as AlphaFold3, which directly predicts the structure of protein–ligand complexes, or by performing conventional docking calculations. Both approaches yield valuable insights into substrate binding and guide the establishment of the metrics that will be used in the filtering criteria to be employed in the platform steps.

Subsequently, candidate enzyme sequences are identified by running BLAST searches of the template sequence against a comprehensive database such as SwissProt (Boutet et al., 2007). For reliable homology modelling, BLAST hits are filtered to retain sequences with at least 30% identity and 60% coverage relative to the template. The resulting list of candidate variants, along with the template structure and the substrate or intermediate structure, are then submitted to the platform’s Gene Discovery feature for ranking and further evaluation.

### Enzyme Engineering

For enzyme engineering applications, it is important to gather information in the literature about the reaction mechanism of each enzyme to help choose the best combination of mutation sites in the protein and the ligand to be used in the process. Whenever possible, experimentally determined structures should be used; otherwise, comparative or *ab initio* models can be generated. With the advent of AlphaFold3 (Abramson et al., 2024), predictions of protein–ligand complex structures are now accessible, providing valuable template models for enzymes and their bound substrate. These approaches yield valuable insights into substrate binding and guide the establishment of the metrics that will be used in the filtering criteria to be employed in the platform steps, as discussed above. The structure of the enzyme to be engineered and the structure of the reaction substrate/intermediate are submitted to the Enzyme Engineering feature of the GDEE platform, as well as the list of residues to be mutated, where enzyme mutants are generated and then evaluated.

### Platform Parameters and User Flexibility

In both protocols, after running the platform pipeline, filters are applied to the results to ensure that ligands are in a catalytically relevant orientation. In the end, a manual inspection is suggested to carefully analyze the results. For the platform parameters, we usually use an optimization radius of 8 Å (matching Vina’s interaction cutoff). As the acceptance thresholds for the modelling step, we follow the threshold proposed by the authors of each method (>0.4 for VoroMQA, < –1 for normalized DOPE). An exhaustiveness value of 200 is typically used unless computational resources are limited (never below 100) and five models are selected for the docking step (see parameter selection section). However, all these parameters and steps can be fully adjusted by the user to accommodate particular needs or specific project requirements.

### Parameters selection

A parameterization process was conducted to find the most suitable values for each method’s parameters. The pipeline parameters were assessed on a data set of over 320 pairs of related proteins, resulting in 10,170 variant models and 305,100 docking calculations. This data set was built by searching on the PDBBind (Liu et al., 2017; R. Wang et al., 2004, 2005) (v2018) for all pairs of related proteins with minor sequence variations. The PDBBind v2018 is a database of X-ray structures of co-crystalized proteins with small compounds for which experimental binding affinity constants are known. The enzyme engineering pipeline was applied to all entries in the aforementioned database with varying algorithms and parameters, in order to evaluate the effects of the individual pipeline steps and their parameters combinations. The most efficient combination of parameters regarding binding affinity sampling and computational processing efficiency were chosen as the default values of the platform.

#### Selection of the docking program and scoring function

First, it was crucial to select the appropriate docking program and scoring function for the pipeline. To this end, we used AutoDock Vina and Smina, each with their respective scoring functions, and Vinardo (Quiroga & Villarreal, 2016) scoring function (within Smina program), to re-dock each ligand’s crystal structure to the corresponding protein.

In enzyme engineering, a precise understanding of how mutations influence ligand binding is essential for designing enzymes with enhanced catalytic properties. With this objective in mind, we analyzed 320 pairs of related proteins to evaluate the accuracy of these methods in predicting the effects of mutations on ligand binding. For each protein pair (reference and variant), we determined the change in binding free energy (ΔΔG) and compared these predictions to experimental data. A method was considered successful (a “Hit”) if it correctly predicted the sign of the free energy difference, i.e., whether the mutation improved or reduced the protein’s binding affinity to the ligand.

Figure 3 presents scatter plots comparing the experimental and predicted ΔΔG values, with percentage of successful predictions referred to as Hits. Among the methods tested, Vina demonstrated the best performance, achieving the highest percentage of successful predictions when considering the pose with the lowest energy or the lowest RMSD for each complex. These results suggest that Vina is a more effective solution for evaluating the enzyme engineering process. However, it is important to note that the ΔΔG values obtained are relatively small, often falling within the typical error margins associated with docking programs. This means that the differences in binding free energy predictions between the reference and variant proteins are subtle and may not be fully captured by the computational methods used. Despite these limitations, the trends observed in our analysis suggest that Vina provides valuable insights, achieving the best performance in the context of enzyme engineering and closely matching Vinardo in previous evaluations. Based on these findings, Vina has been selected as the default scoring function in our platform.

**Figure 3.**
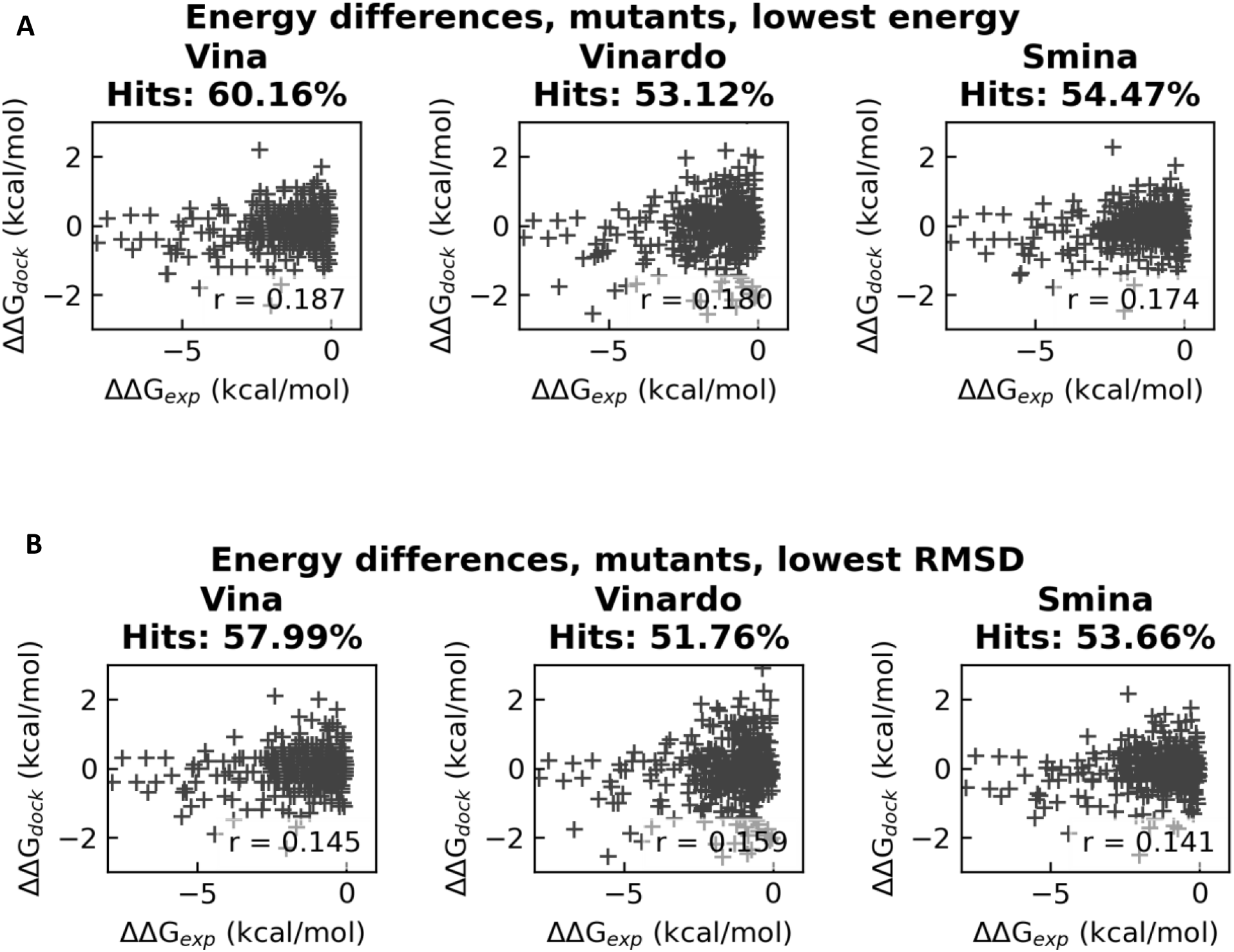
Differences in binding affinities of mutant pairs from docking calculations and experimental data. Hits represent the percentage of docking-based calculations that were able to qualitatively predict the changes in binding affinities from experimental data. For each protein, in (A) we considered the docking poses with the lowest energy and in (B) the docking poses with the lowest RMSD (B).

#### Selection of exhaustiveness and number of models

After selecting the docking software and scoring function, it was necessary to determine the appropriate values of exhaustiveness and the number of models used in the docking step. Figure 4 presents heatmaps showing the energy distribution of docking solutions across the tested parameter space (exhaustiveness and number of models), corresponding to different numbers of models used. Each heatmap column represents the energy distribution for a fixed exhaustiveness value. In these heatmaps, a reduction in variance is observed when increasing the number of models from one to five. However, beyond five models, no significant further reduction is observed, indicating that statistical sampling is effectively achieved at five models.

**Figure 4.**
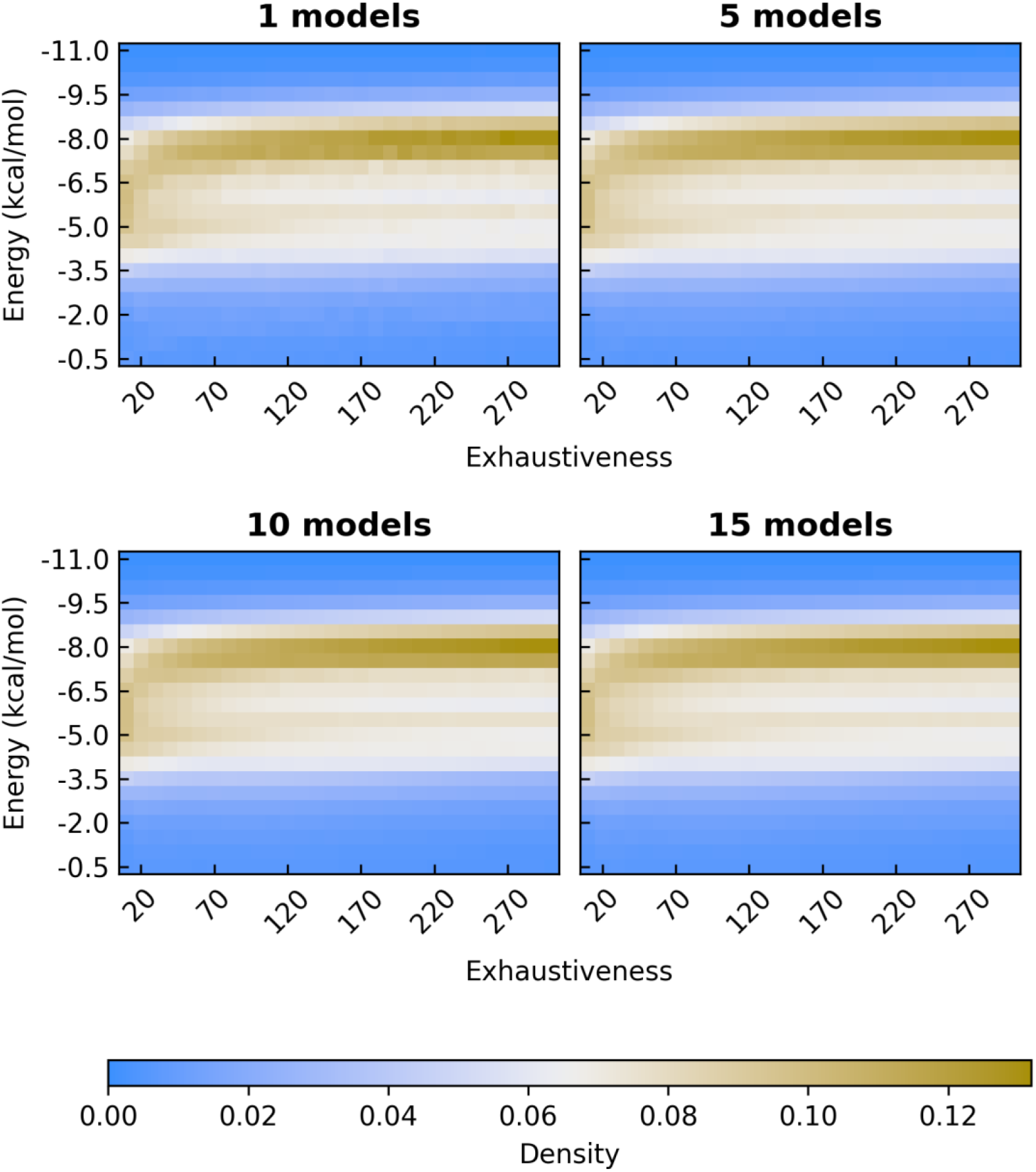
Heatmaps showing the energy distribution of docking solutions for varying number of models used across different exhaustiveness values. Each heatmap column represents the energy distribution for a fixed exhaustiveness value. Color intensity within each column indicates the density of energy values, with higher density represented by dark golden yellow. Noticeable, differences in energy distribution and reduced noise are observed when the number of models increases from one to five. Beyond five models, further increases do not result in visible improvements.

This convergence of variance with multiple models is further supported by the data in Figure 5 (bottom plot), reinforcing the conclusion that five models provide the right balance between comprehensive conformational sampling and computational efficiency.

**Figure 5.**
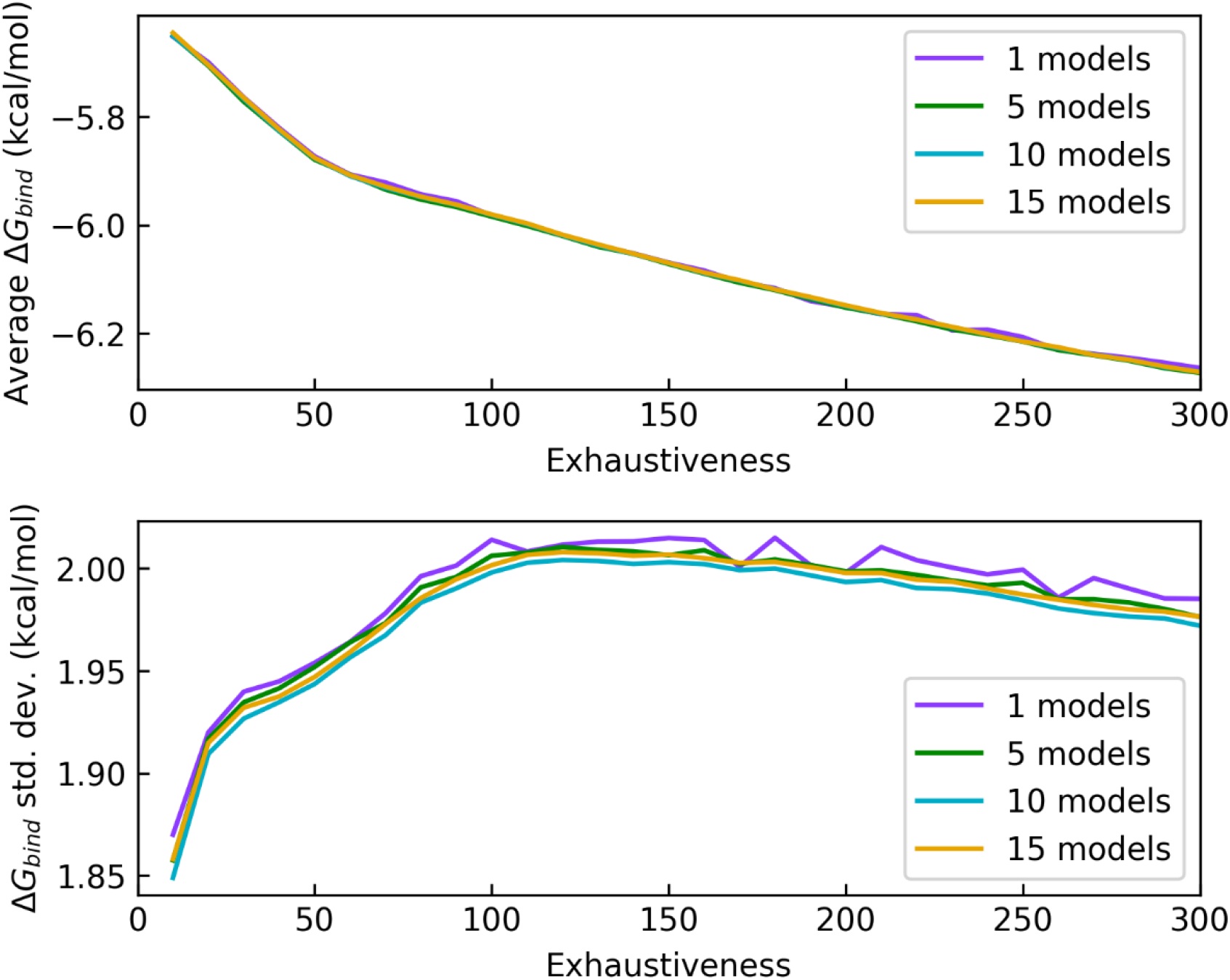
Comparison of average binding affinity (top plot) and standard deviation of binding affinities (bottom plot) across exhaustiveness values for different number of models used. The bottom plot, clearly demonstrates a smoother standard deviation curve when using more than one model, compared to results from a single model, indicating improved sampling and lower noise.

Figure 5 complement these findings, showing that increasing the exhaustiveness parameter from 100 to 300 yields only a minor decrease (∼0.2 kcal/mol) in average binding energy. This suggests the docking process has essentially converged within this exhaustiveness range.

Based on these results, we recommend using five models and an exhaustiveness of at least 100 as the parameters for docking steps in the platform. These settings provide an effective compromise between accuracy and computational cost and have been adopted as the standard configuration in our protocols.

### Re-scoring of docking poses with machine learning scoring functions

The platform has been applied to multiple case studies, with several successes (two of which are already published (Ferreira et al., 2024; Hanko et al., 2023), with others forthcoming) as well as a few less successful outcomes. These unsuccessful cases highlighted areas for improvement, particularly in the platform’s scoring accuracy, which is crucial for enhancing its ranking capabilities.

The docking step is crucial to evaluate the variants generated by the platform. Despite current algorithms managing pose generation effectively, the imprecision of current scoring methodologies is the main obstacle to achieving reliability in docking. To reduce this limitation, a re-scoring methodology will be added to the platform. This method utilizes machine learning-based (ML) scoring functions to re-score the poses obtained by the docking step. These non-parametric machine learning methods can be used to implicitly capture the binding interactions that are challenging to model explicitly. By not imposing a particular functional form for the scoring function, the collective effect of intermolecular interactions in binding can be directly inferred from experimental data (Ballester & Mitchell, 2010; Durrant & McCammon, 2010; Pereira et al., 2016). This method has already been shown to improve scoring performance, as demonstrated by several authors (Ballester et al., 2014; Ballester & Mitchell, 2010; Durrant & McCammon, 2010; Li et al., 2015; Pereira et al., 2016). For this task, seven machine learning-based scoring functions (Rf-Score v1 (Ballester & Mitchell, 2010), Rf-Score v2 (Ballester et al., 2014), Rf-Score v3 (Li et al., 2015), NNScore (Durrant & McCammon, 2010, 2011), PLECrf, PLECnn and PLEClinear (Wójcikowski et al., 2019)) were selected from the literature and chosen for further testing and comparison with classical ones (Vina, Smina, Vinardo and AutoDock4.2 (AD4) (Morris et al., 2009)).

#### Scoring precision

First, all scoring functions were evaluated based on their scoring precision using the pose of 4010 protein-ligand complexes from the PDBBind v2018 dataset (See Figure 6) (All scatter plots from the re-scoring study are provided in Figures S1 to S9 of the supplementary materials). Scoring function of Vinardo exhibited the lowest Mean Squared Error (MSE), but its coefficient of determination (R^2^) was lower than that of several machine learning-based models. Scoring functions of Vina, Smina, and AD4 showed similar results, with comparable MSE and R^2^ values.

**Figure 6.**
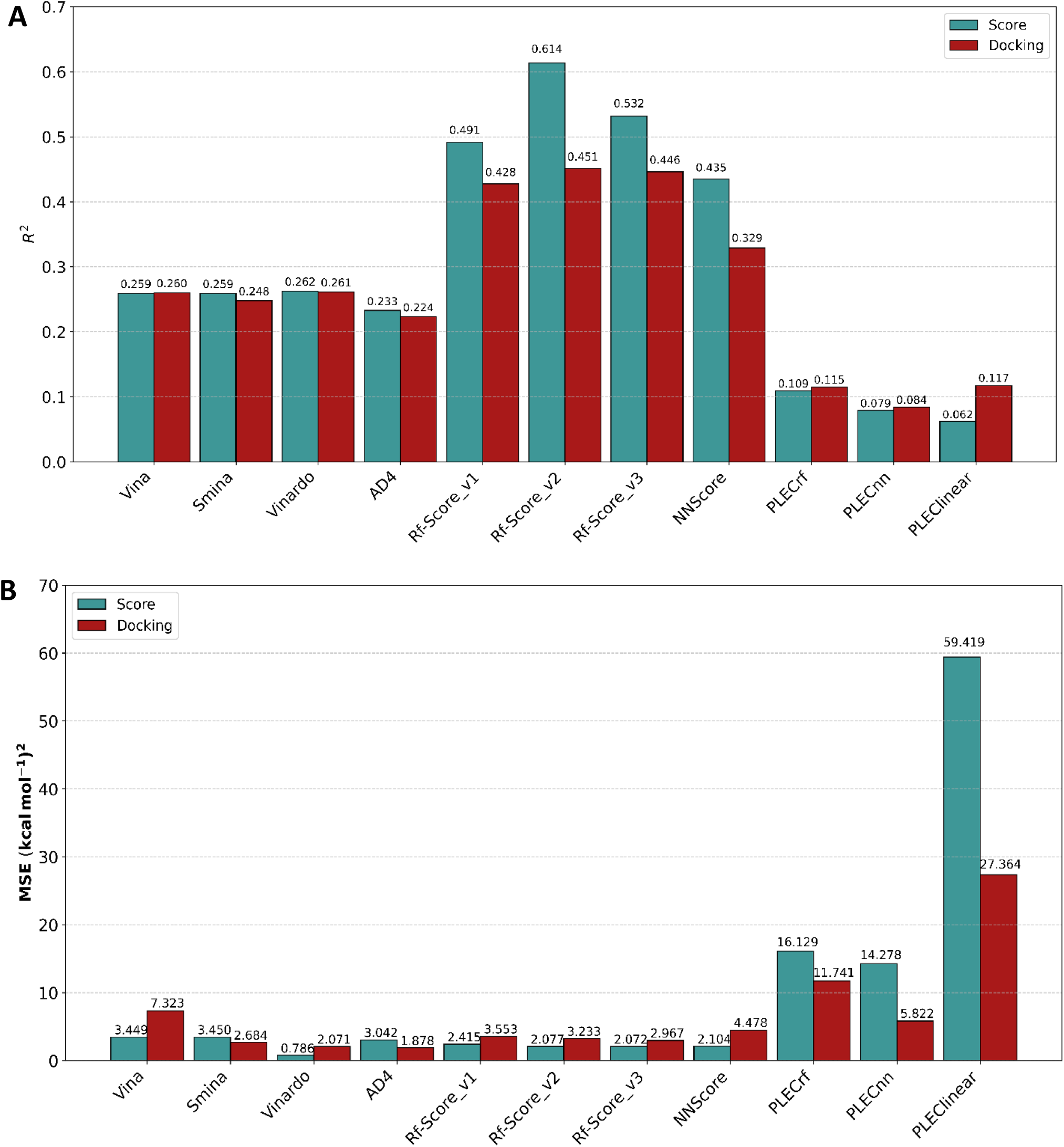
R_2_ (A) and MSE (B) of tested scoring functions evaluated on 4010 protein-ligand complexes from the PDBBind v2018 dataset. The teal bars correspond to performance on X-ray structures, while the red bar indicate results obtained after re-docking the ligands into protein binding pockets using AutoDock Vina, considering only the lowest-energy pose identified by Vina. The Vinardo scoring function exhibited the lowest MSE on X-ray structures but had a lower R_2_ compared to several machine learning-based models. Traditional scoring functions such as Vina, Smina, and AD4 demonstrated comparable MSE and R_2_ values. ML models generally improved predictive accuracy, except for PLEC Score functions, which showed poor performance with high MSE and low correlation. After docking, Vina and Vinardo’s performance decreased in MSE, while AD4 and Smina showed slight improvements. Machine learning models maintained consistent correlation levels, with Rf-Score v2 achieving the highest R_2_ value.

Most machine learning models improved performance, except for the PLEC Score versions. Rf-Score v2 was the best performer overall, achieving the highest R^2^ and one of the lowest MSEs. In contrast, the PLEC Score models had poor results, with significantly higher MSE and weak correlation.

Figure 6 also includes an evaluation of the scoring functions’ accuracy following docking calculations, with the ligands being re-docked into protein binding pockets using AutoDock Vina. The docking results were re-scored using AD4, Smina, Vinardo, and the ML models, with only the lowest energy pose found by Vina being considered.

The results indicated that Vina and Vinardo performed worse in terms of MSE after docking, while AD4 and Smina showed slight improvements. The four ML scoring functions (excluding PLEC versions) maintained similar correlation levels as in the initial evaluation. The ML models performed consistently with Rf-Score v2 once again achieving the highest R^2^. PLEC Score models consistently underperformed in all evaluations, yielding the poorest overall results. Therefore, they have been omitted from the subsequent figures, but their detailed results are available in the supplementary materials.

#### Docking precision

Docking precision refers to the ability of a scoring function to select a ligand pose that closely matches the crystal structure as the best pose from a set of poses generated during docking. This was assessed by scoring all poses from the docking process and determining whether the scoring function could identify a pose with an RMSD within 2 Å of the crystal structure as the most favorable in terms of predicted affinity. The results are presented in Figure 7.

**Figure 7.**
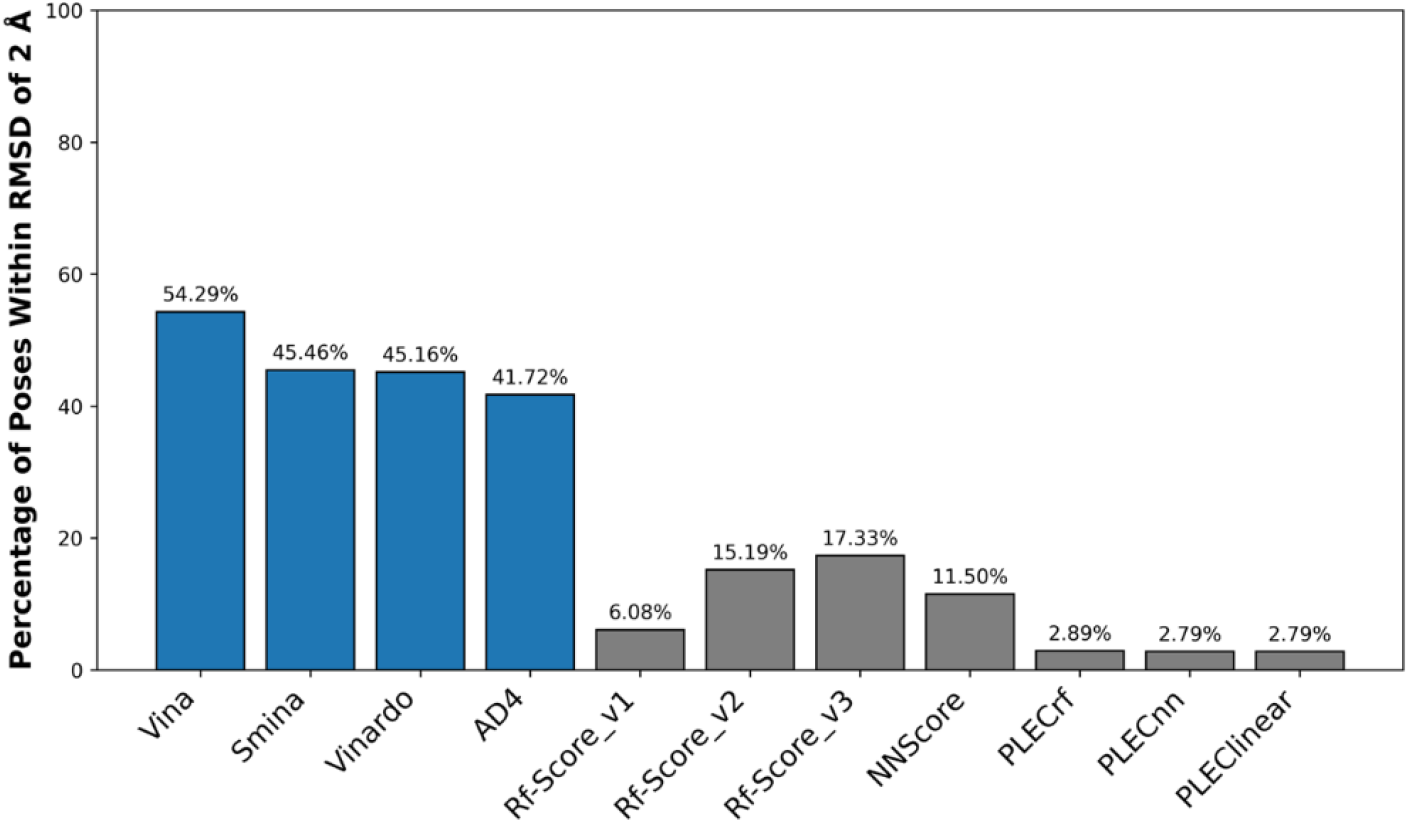
Docking power evaluation of scoring functions on 4010 protein-ligand complexes. Docking power was measured as the percentage of predicted poses within 2 Å RMSD of the corresponding X-ray structure, among the lowest-energy poses identified by each scoring function. Classical scoring functions are shown in blue and ML-based scoring functions in grey. Among all tested methods, Vina achieved the highest docking power, outperforming all ML models. The generally lower performance of ML scoring functions in docking power may be attributed to their training primarily on experimented determined complexes rather than on docked poses, which limits their ability to accurately identify correct bound conformations.

Classical scoring functions significantly outperformed ML models in docking precision. The scoring function of Vina was the most effective, correctly identifying 54% of poses within 2 Å of the crystal structure. Smina, Vinardo and AD4 followed with around 45% accuracy each. In contrast, machine learning models performed poorly, with accuracy ranging from just 2% to 17%. Among them, versions 2 and 3 of Rf-Score and NNScore showed the best results, though still well below the classical functions.

These findings suggest that machine learning-based scoring functions are less suited for selecting the best poses from docking results. Their strength lies primarily in improving the accuracy of predicted binding affinities for correctly docked poses. This limitation may stem from their training on databases of experimentally determined structures, which allows them to learn correct poses but not effectively distinguish incorrect ones.

#### Evaluating Scoring Within the Scope of Enzyme Engineering

To assess the accuracy of the scoring functions in enzyme engineering, the previous results were organized into 268 pairs of related proteins to evaluate how well the functions predicted the effects of mutations on binding.

Figure 8 depicts the percentage of successes, of each scoring function, on predicting the mutation’s effect on binding. Success is defined as correctly predicting the sign of the ΔΔG. Scoring functions of Vina, Smina, Vinardo, and AD4 correctly predicted approximately 50% of cases, both from X-ray structures and after docking calculations. Rf-Score v2 had the best performance, with a success rate of 68% on X-ray structures and 60% after docking. NNScore also demonstrated an improved success rate, around 60% in both scenarios. The remaining models performed similarly or worse than the classical scoring functions.

**Figure 8.**
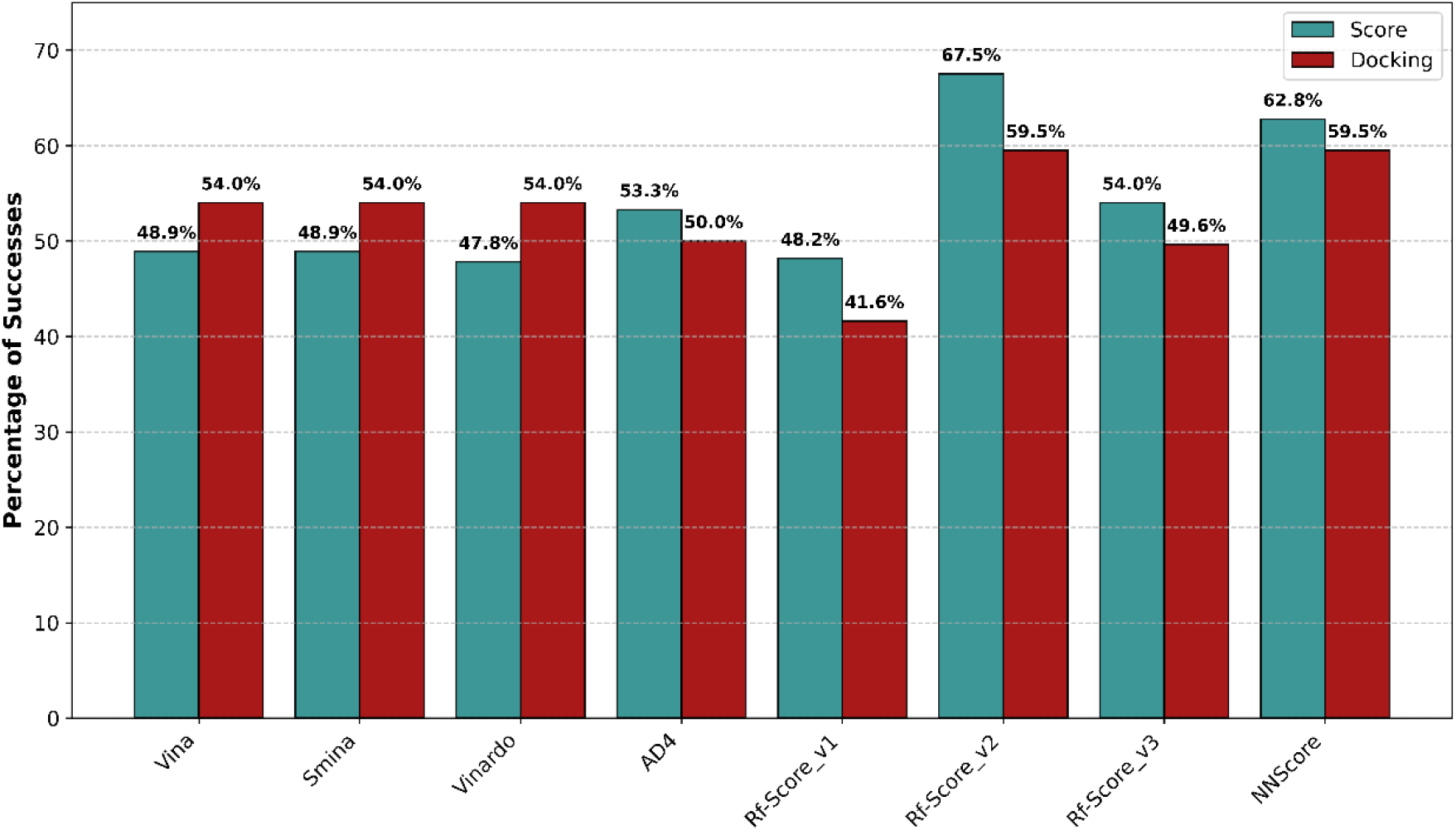
Percentage of successes for each scoring function in predicting the mutation’s effect on binding affinity within 268 of reference-variant pairs. Success is defined as the fraction of cases in which the scoring function correctly predicted the sign of the change in binding free energy (ΔΔG) compared to experimental data. Results are shown for scores calculated using both X-ray crystal structures (teal) and structures after ligand re-docking into the protein’s active site (red), based on the lowest-energy pose identified by Vina. Classical scoring functions including Vina, Smina, Vinardo, and AD4 achieved approximately 50% success rates across both structural contexts. Among machine learning models, Rf-Score v2 performed best, with success rates of 68% on X-ray structures and 60% after docking, while NNScore also showed improved rates near 60%. Other ML models displayed similar or reduced accuracy compared to classical functions. This assessment underscores the variations in predictive accuracy introduced by mutation effects and structural modeling, informing the selection of the best scoring approaches for enzyme engineering applications.

### Stacking Ensemble of Scoring models

Building on our finding that certain machine learning-based scoring functions improve binding affinity predictions, we explored whether combining these models into an ensemble could further enhance performance. We employed stacking generalization as the ensemble method, integrating the scoring functions studied earlier to create a metamodel in an attempt to enhance scoring performance of the GDEE platform. We implemented linear regression as the stacking model due to its interpretability and effectiveness in combining diverse model outputs.

To train the stacking model, 75% of the dataset was used, with the remaining 25% reserved for validation. This validation set includes the 268 pairs of reference and mutant proteins, but also contains additional samples beyond these pairs, ensuring that the model could be tested in the context of enzyme engineering. By using the same subset across all scoring functions, a fair comparison of performance was achieved.

#### Exploring all model combinations

To optimize the metamodel, we conducted a comprehensive evaluation of all possible combinations of re-scoring models, including Vina, Smina, Vinardo, AD4 and all ML models. The evaluation was based on two metrics: MSE and correlation coefficient (R^2^). This approach aimed to identify the model ensemble that would best enhance predictive accuracy, providing key insights into selecting base models for the stacking process.

Figure 9 illustrates the correlation between the number of models and performance metrics (MSE and R^2^). Each data point represents the metric values achieved for the best combination of a given number of models. The results demonstrate that increasing the number of models from one to five yields improvements in both metrics. However, beyond five models, further increases result in only small changes in the performance metrics, indicating that additional models offer minimal benefit. For efficiency, using more than five models is not justified, as the gains in predictive performance become limited.

**Figure 9.**
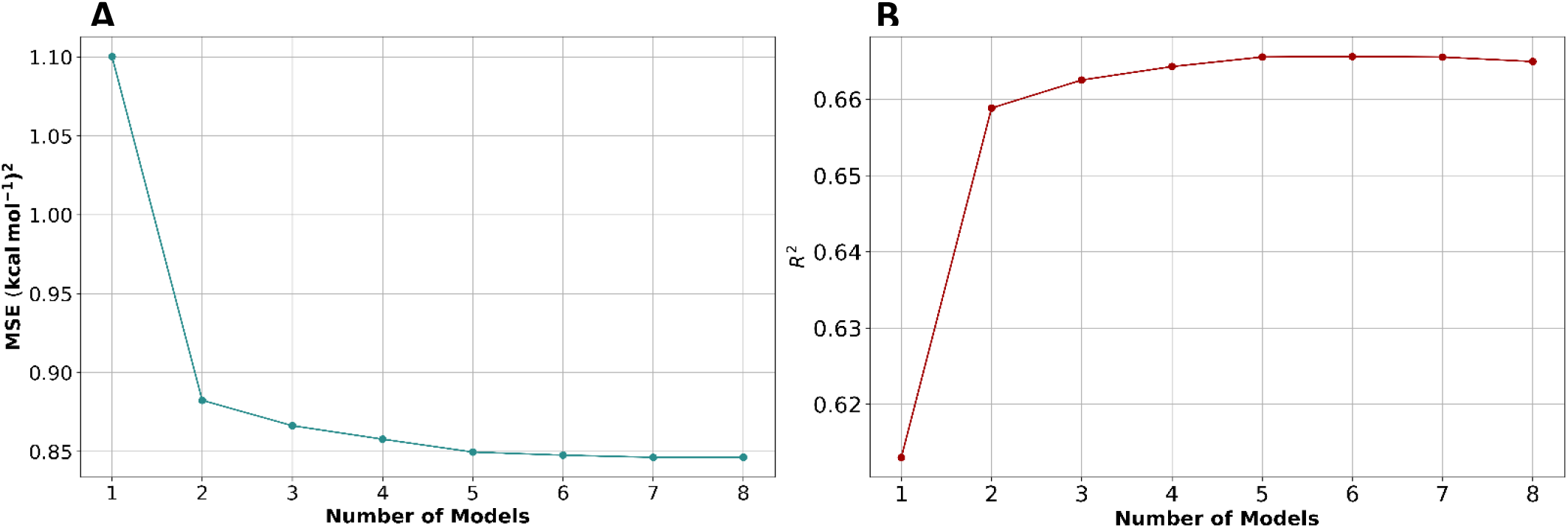
Relationship between the number of base models used in the stacking ensemble and predictive performance metrics. Plotted are the best achieved MSE (A) and R_2_ (B) for each model combination size, evaluated on the validation set. Each data point corresponds to the best combination of base models yielding the best performance for that given number of models. The results reveal a significant improvement in both MSE and R_2_ as the number of models increases from one to five, demonstrating the benefits of ensemble methods in reducing prediction errors. Beyond five models, marginal gains in performance become negligible, suggesting minimal returns with the inclusion of additional base models. Consequently, limiting the ensemble to five base models strikes an effective balance between predictive accuracy and computational efficiency.

The best combination, consisting of five base models, is detailed in Table 1, along with their respective weights. Notably, Rf-Score v2 carries significantly more weight than the others, reflecting its superior individual performance. This metamodel was then applied to re-score protein-ligand complexes, and its performance was compared against other models.

**Table 1.** Composition and relative weights of the five base models in the optimized stacking ensemble metamodel. This table summarizes the individual base models selected for the ensemble and their assigned weights reflecting their contribution to the final predictive model. Notably, Rf-Score v2 carries significantly greater weight than other models, consistent with its superior standalone performance. The weighted combination effectively integrates diverse predictive strengths and was subsequently applied to re-score protein-ligand complexes, demonstrating enhanced accuracy compared to individual scoring functions.

| Base model | Weight |
| --- | --- |
| $\beta_0$ | 3.439 |
| Rf-Score v1 | -0.976 |
| Rf-Score v2 | 2.640 |
| Rf-Score v3 | -0.282 |
| PLECnn | 0.060 |
| Vina | -0.044 |

Figure 10 presents histograms comparing predicted and experimental binding affinities, both for the X-ray structures and after re-docking ligands using AutoDock Vina. The metamodel outperformed all individual scoring functions. While Vinardo had a comparable MSE (see Figure S6 in supplementary materials), the metamodel significantly outperformed it in R^2^, indicating its superior ability to explain the variance in the data.

**Figure 10.**
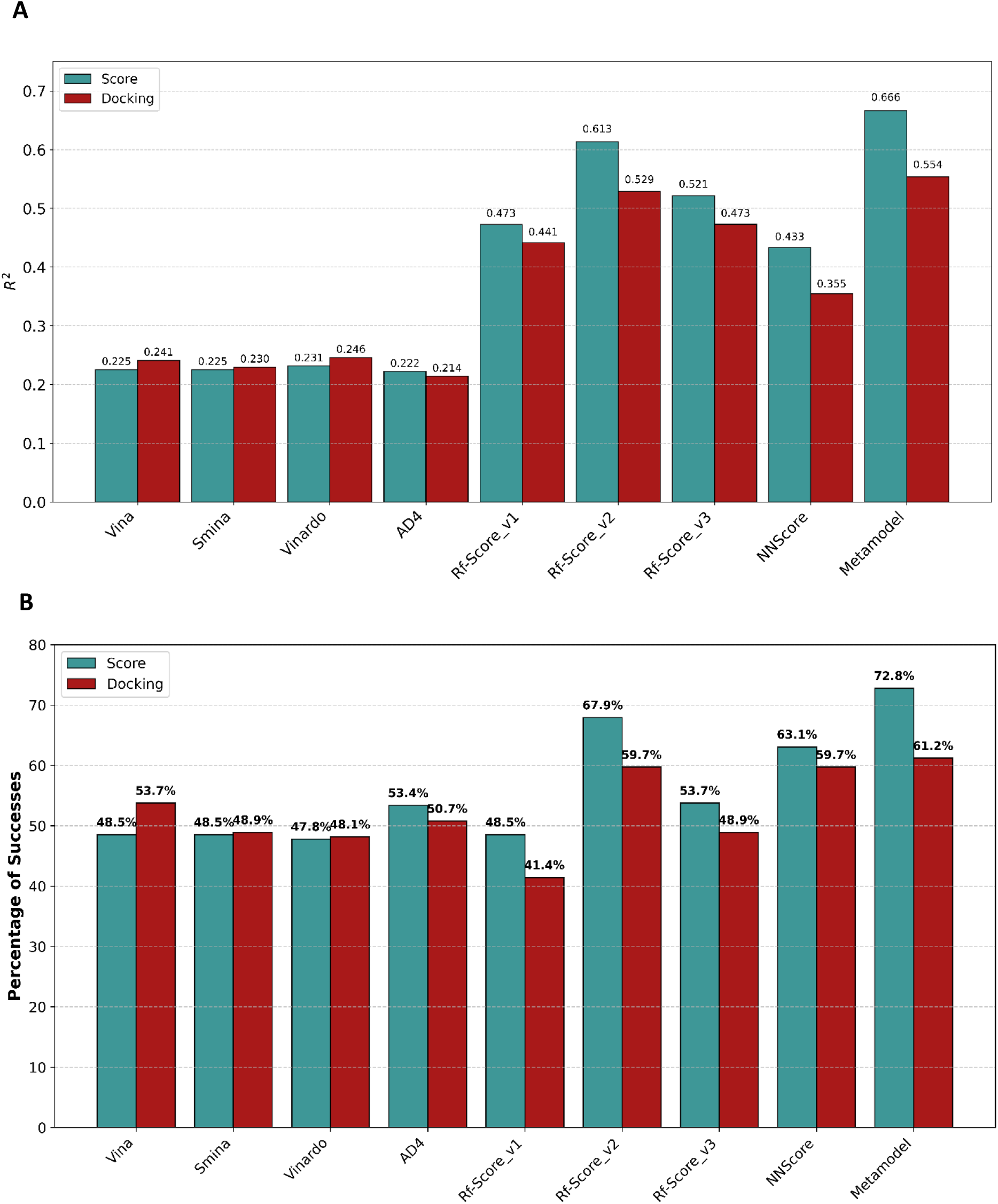
Histogram comparison of predicted versus experimental binding affinity metrics for the stacking validation set. (A) R_2_between predicted and experimental values for X-ray crystal structures (teal) and ligand re-docked structures using AutoDock Vina (red), based on the lowest-energy pose identified by Vina. (B) Percentage of successes in predicting the sign of the mutation’s effect on binding affinity (ΔΔG) for the same structural contexts. The metamodel outperformed all individual scoring functions, exhibiting superior R_2_ despite Vinardo showing comparable MSE (see Figure S6 in the supplementary materials). For mutation subsets within the database, the metamodel improved success rates beyond the best individual model (Rf-Score v2), reaching 73% on X-ray structures and 61% post-docking. This underscores the metamodel’s ability to effectively combine information from multiple scoring functions to enhance mutation impact predictions.

When analysing the mutations within the database (Figure 10B), the metamodel again showed superior performance, improving Rf-Score v2’s success rate (the best individual function) to 73% and 61% for the respective cases. This represents an improvement over Rf-Score v2 alone, highlighting the metamodel’s effectiveness in combining information from multiple scoring functions to predict the impact of mutations on binding affinity.

While comparing the Gibbs free energy of interaction between reference and variant enzymes, calculated from X-ray structures, offers valuable insights for enzyme engineering, this work focuses specifically on evaluating scoring functions within the broader context of the platform’s pipeline. This distinction is important because the platform’s workflow encompasses not only docking procedures but also the prediction of mutant enzyme structures (steps that were not included in the previous analysis). Thus, the current evaluation captures the integrated performance of scoring functions as applied to predicted mutant structures, offering a more comprehensive assessment beyond direct experimental structures.

To conduct this evaluation, we proceed on applying the platform to the 167 reference proteins, independent proteins that have mutants in the database. These 167 reference proteins, corresponding to the 268 pairs of reference-variant proteins (as some references have multiple variant counterparts), were input into the platform to generate structural models, perform docking calculation with the respective ligand and calculate binding affinities for their respective variants. All scoring functions, except for Vina, which was already integrated into the platform, were then used to re-score the resulting complexes.

This approach provides a comprehensive assessment of the scoring functions across the platform’s entire pipeline, offering insights into their performance beyond individual docking procedures. However, it is important to note that this evaluation involved fewer complexes compared to the previous analysis shown in Figure 6, because not every protein in the dataset has a corresponding variant. As a result, direct comparisons between the two evaluations are not feasible.

Figure 11 shows the comparison between experimental and predicted binding affinity values for all complexes, both reference and variant, after re-scoring the best pose identified by Vina. As previously observed, all three versions of Rf-Score continued to provide high predictive performance regarding *R*^*2*^, surpassing the other scoring functions. However, the metamodel achieved superior results in terms of MSE, delivering the lowest observed error among all methods.

**Figure 11.**
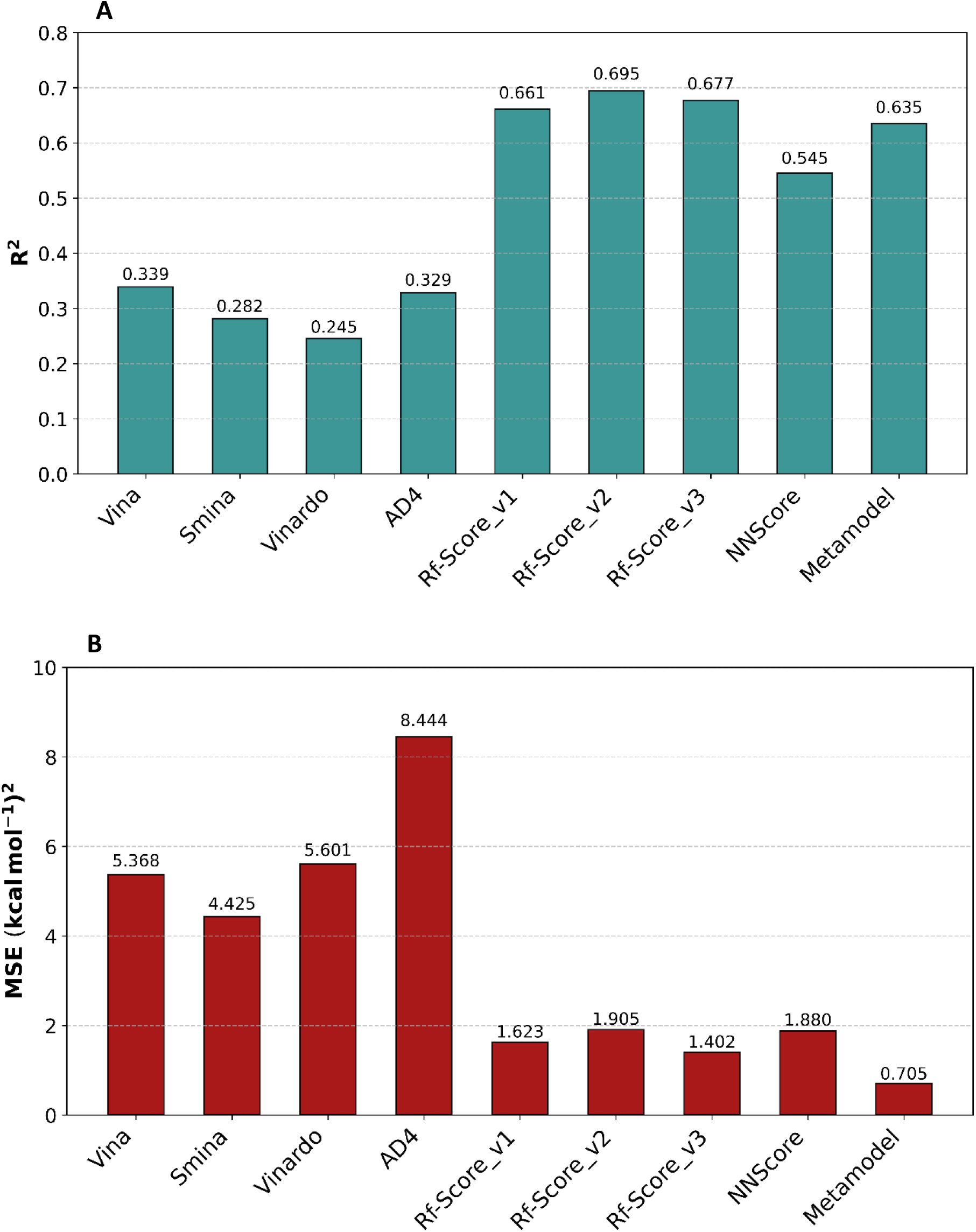
Histogram comparison of R_2_ (A) and MSE (B) between predicted and experimental binding affinities obtained through the platform’s pipeline in the stacking validation set. The results are based on the lowest-energy pose identified by Vina. Among the scoring functions evaluated, the three versions of Rf-Score achieved the highest R_2_ performance, closely followed by the metamodel. For the MSE metric, the metamodel surpassed all other scoring functions, indicating superior accuracy in predicting binding affinity. These results highlight the effectiveness of the metamodel in integrating diverse predictive signals to enhance overall performance

Figure 12 presents the percentage of successes for each scoring function, defined as the fraction of cases where the sign of ΔΔG was correctly predicted for each mutation. Vinardo demonstrated the highest success rate, with correct predictions in nearly 55% of cases for the 268 pairs of reference and variant proteins. In contrast, most other scoring functions exhibited a decrease in success rates compared to previous phases (Figure 10), which may result from uncertainties introduced by the protein structure prediction methodology. An additional consideration is that the protein models were constructed (by Modeller) without the ligand present in the enzyme’s active site, potentially introducing subtle changes in protein conformation and consequently affecting binding predictions.

**Figure 12.**
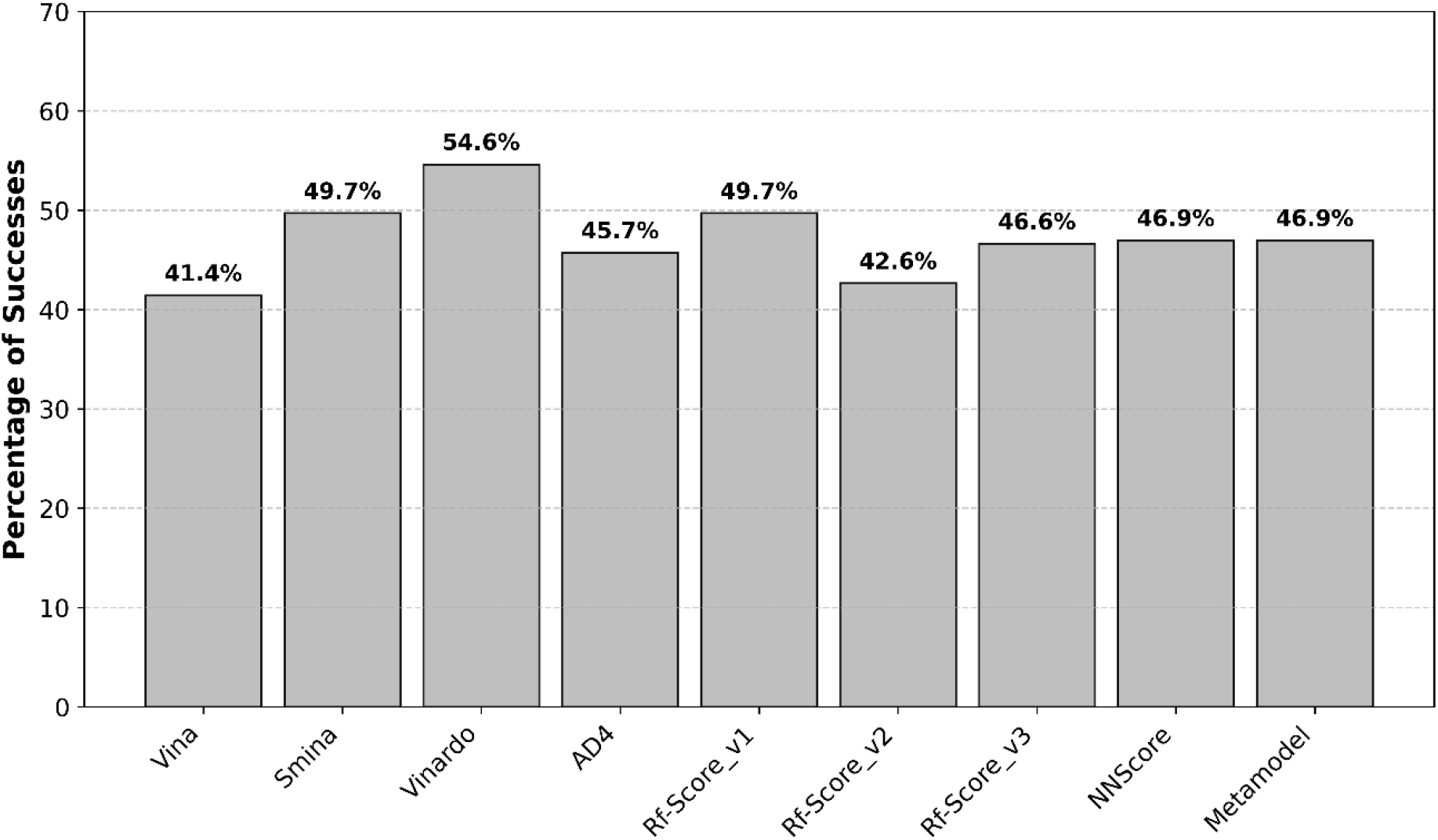
Percentage of successes for predicting mutation effects on binding affinity across scoring functions using the platform’s pipeline in the stacking validation set. Success is defined as correctly predicting the qualitative change in binding affinity (i.e., the sign of ΔΔG) based on experimental measurements. Predictions consider the lowest-energy pose obtained by Vina. The histogram illustrates the comparative performance of multiple scoring functions on the 268 reference-variant pairs, highlighting variations in predictive accuracy. Vinardo achieved the highest success rate, while other scoring functions exhibited diverse results, potentially influenced by structural modelling uncertainties during protein structure prediction.

Based on these results, the metamodel was selected as the preferred scoring function for re-scoring docking poses due to its strong and consistent performance across distinct evaluation scenarios. In the initial benchmarking, performed independently of the platform’s full pipeline, the metamodel achieved the highest predictive accuracy, outperforming all individual scoring functions on all performance metrics and demonstrating overall reliability. This findings are consistent with broader research in protein-ligand binding affinity prediction, where ensemble and meta-modeling approaches have shown greater robustness and generalizability than single models by integrating complementary predictive signals (Lee et al., 2024; Mohamed Abdul Cader et al., 2024).

Subsequent evaluation within the platform’s pipeline further confirmed the metamodel’s value. Although it did not always achieve the absolute top score in every metric, it maintained highly competitive results, including an MSE of 0.7 (the lowest among all methods tested), an *R*^*2*^ of 0.63 (close to the best at 0.69), and a success rate of 46.9% (with the top at 54%). The ability of the metamodel to sustain strong performance, even when faced with structural uncertainties and data complexity introduced by the platform pipeline, underscores its resilience and flexibility in practical, integrative applications.

Taken together, both independent and platform-integrated evaluations highlight the metamodel as the most beneficial choice for the platform’s re-scoring step. Its capacity to deliver high accuracy and reliable, high-quality predictions in realistic operational conditions makes it a suitable solution for protein-ligand binding affinity predictions.

## Conclusions

Enzyme engineering has become a cornerstone in numerous fields, ranging from pharmaceuticals and biocatalysis to sustainable energy and environmental remediation. The growing demand for custom-designed enzymes with enhanced stability, specificity, and catalytic efficiency underscores the need for advanced tools that can accelerate and optimize the engineering process. In this work, we introduce a comprehensive computational platform specifically designed to address this need, offering a powerful suite of tools for identifying natural enzymes capable of catalyzing specific reactions and to facilitate the *in-silico* optimization of enzyme properties.

The case studies referenced here exemplify the GDEE platform’s potential for broad applications in synthetic biology, rapidly identifying and optimizing enzyme variants for specific functions, demonstrating its potential as a powerful tool in metabolic engineering.

Additionally, this study highlights the potential of ML scoring functions to significantly enhance the accuracy of binding affinity predictions in protein-ligand complexes. While these models show great promise in predicting binding affinities, their effectiveness in selecting the optimal pose from docking calculations is limited. This limitation likely stems from their training on experimentally determined structural data, which enables them to learn correct poses but restricts their ability to recognize incorrect ones.

Given these insights, the most effective protocol involves performing docking calculations using AutoDock Vina, which demonstrated superior accuracy in identifying the best pose. Then, re-scoring this pose with a ML scoring function improves the predictive accuracy of binding affinity. This combined approach leverages Vina’s strength in pose selection and the precision of machine learning models for affinity prediction.

To incorporate these findings, re-scoring with the metamodel developed here, it will be implemented as an optional step in the GDEE platform after the filtering phase, allowing users to re-score the poses identified by Vina and thereby improve ranking precision.

## Availability

The source code of GDEE platform will be available in https://github.com/protein-modelling-itqb/gdee.

