## Supplementary material for "GDEE: A Structure-Based Platform for Gene Discovery and Enzyme Engineering"

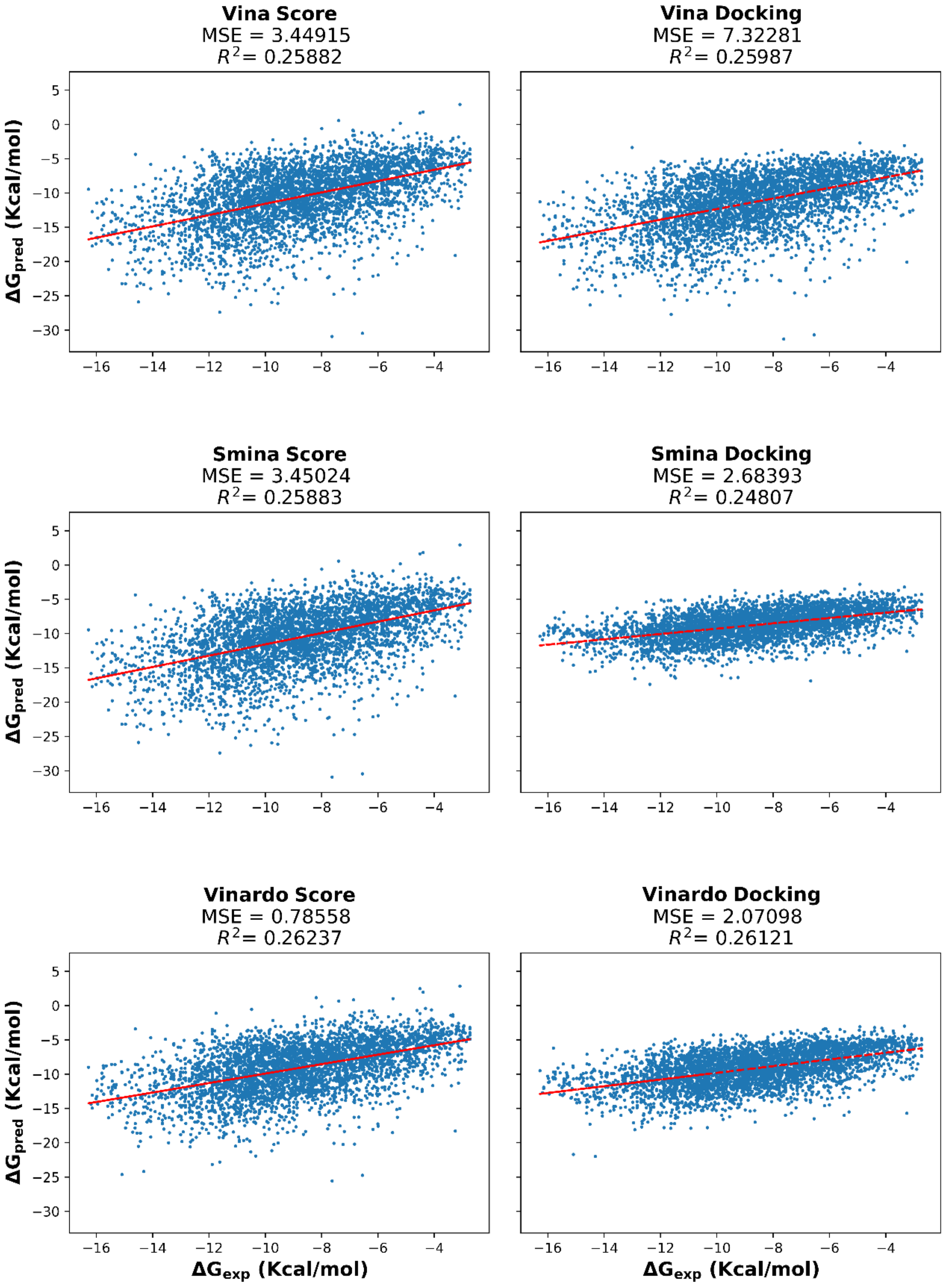


*Figure S1 - Scatter plots comparing predicted binding affinities to experimental binding affinities. Left column - Predicted binding affinities when calculated using each scoring function on the crystallized pose (first phase). Right column – predicted binding affinities after re-docking the ligand in the protein's binding pocket (second phase). The result of a linear regression is represented in red. Structures from PDBbind v2018 database. (Page 1 of 4).*

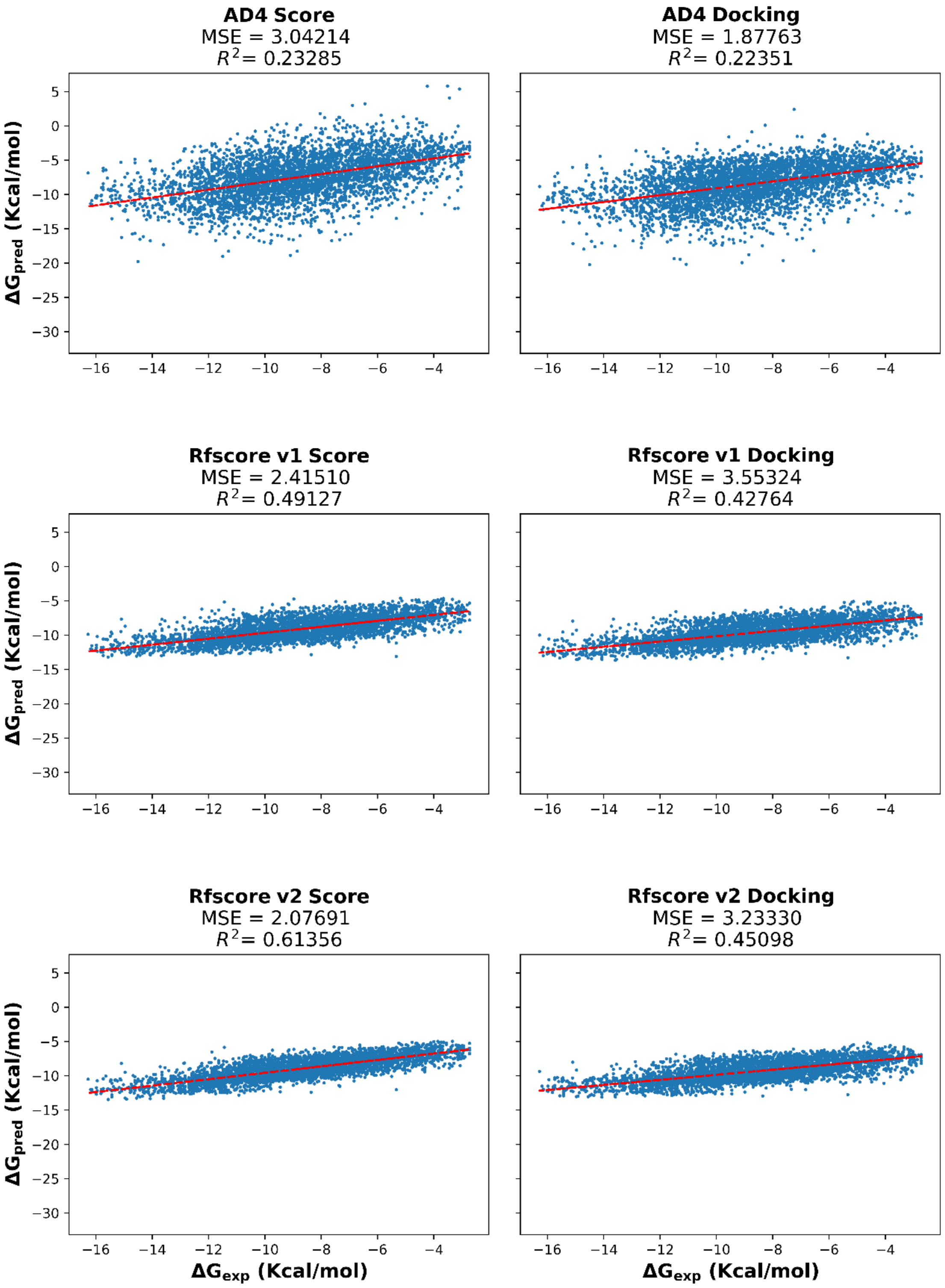


*Figure S1 - Continued (Page 2 of 4).*

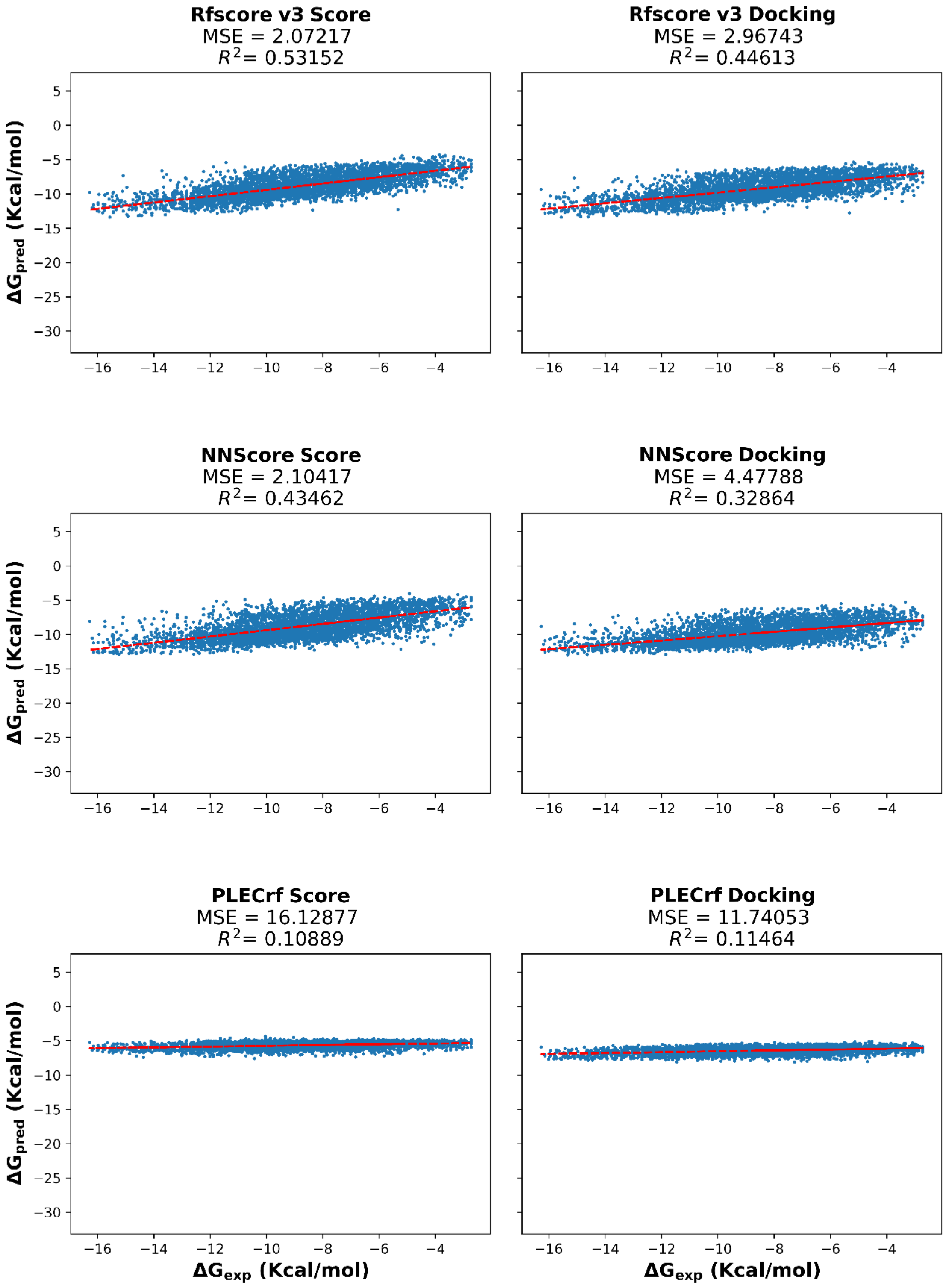


*Figure S1 - Continued (Page 3 of 4).*

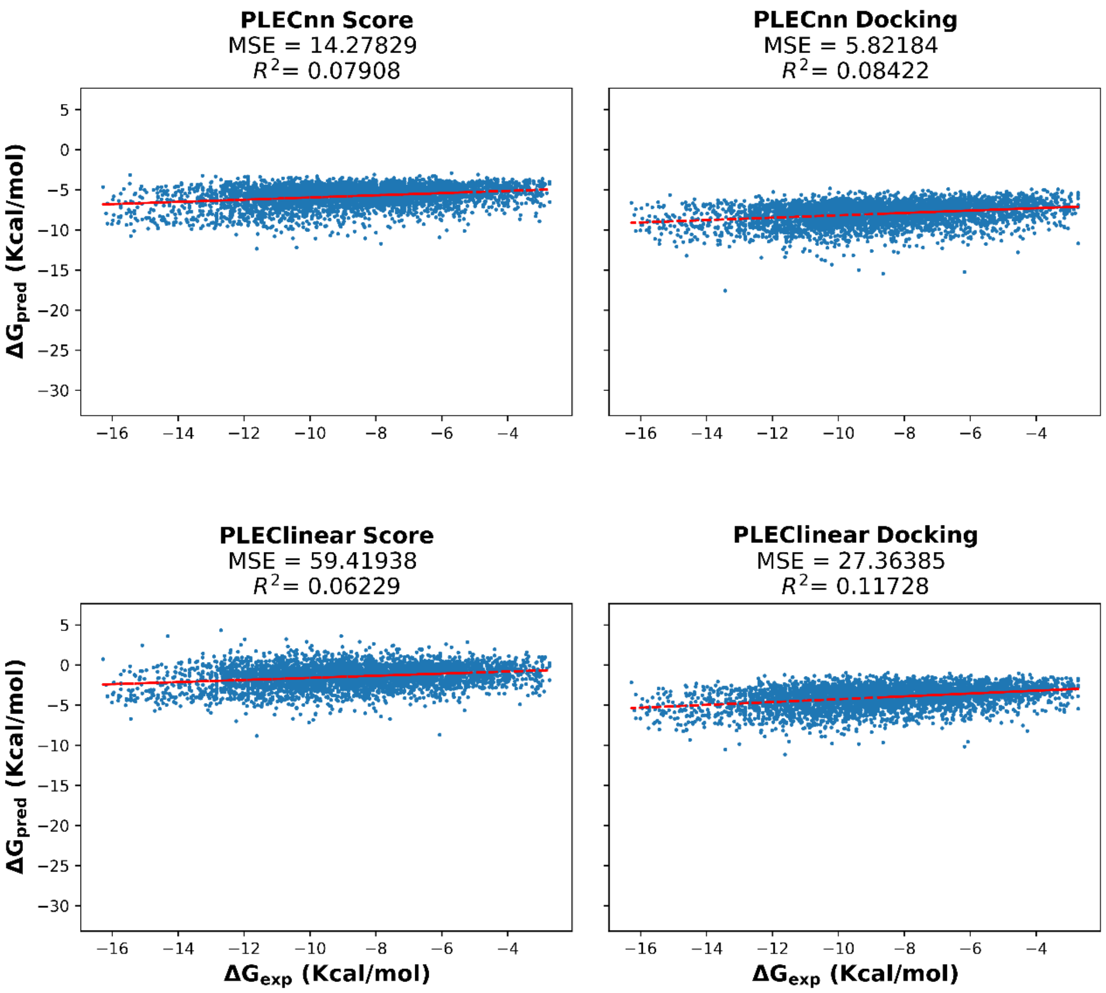


*Figure S1 - Continued (Page 4 of 4).*

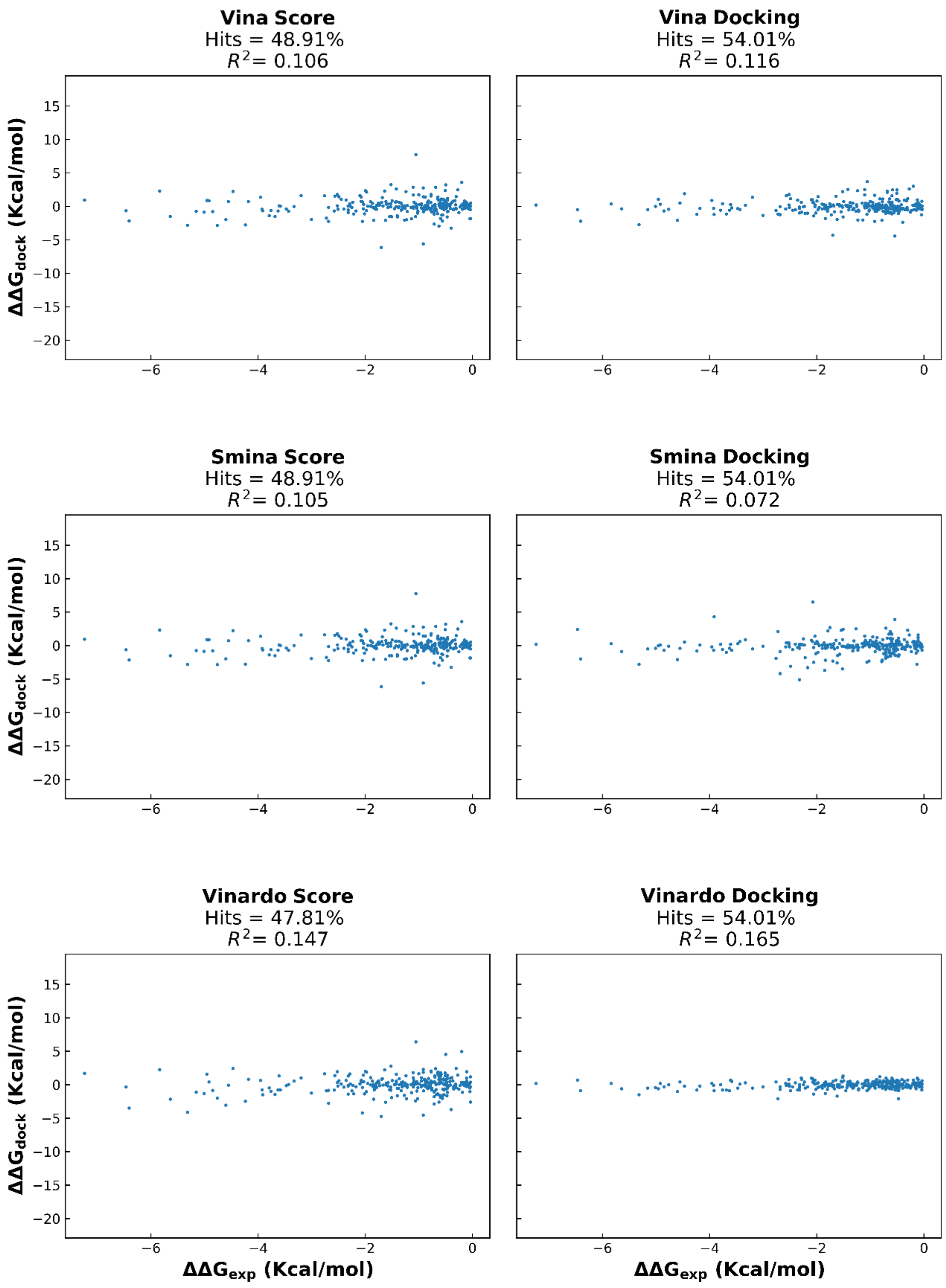


*Figure S2 - Scatter plots comparing predicted ΔΔG to experimental ΔΔG. Left column - Predicted ΔΔG when calculated using each scoring function on the crystallized pose (first phase). Right column – predicted ΔΔG after re-docking the ligand in the protein's binding pocket (second phase). Hits – Percentage of successes on predicting the mutation's effect on binding. Structures from PDBbind v2018 database. (Page 1 of 4).*

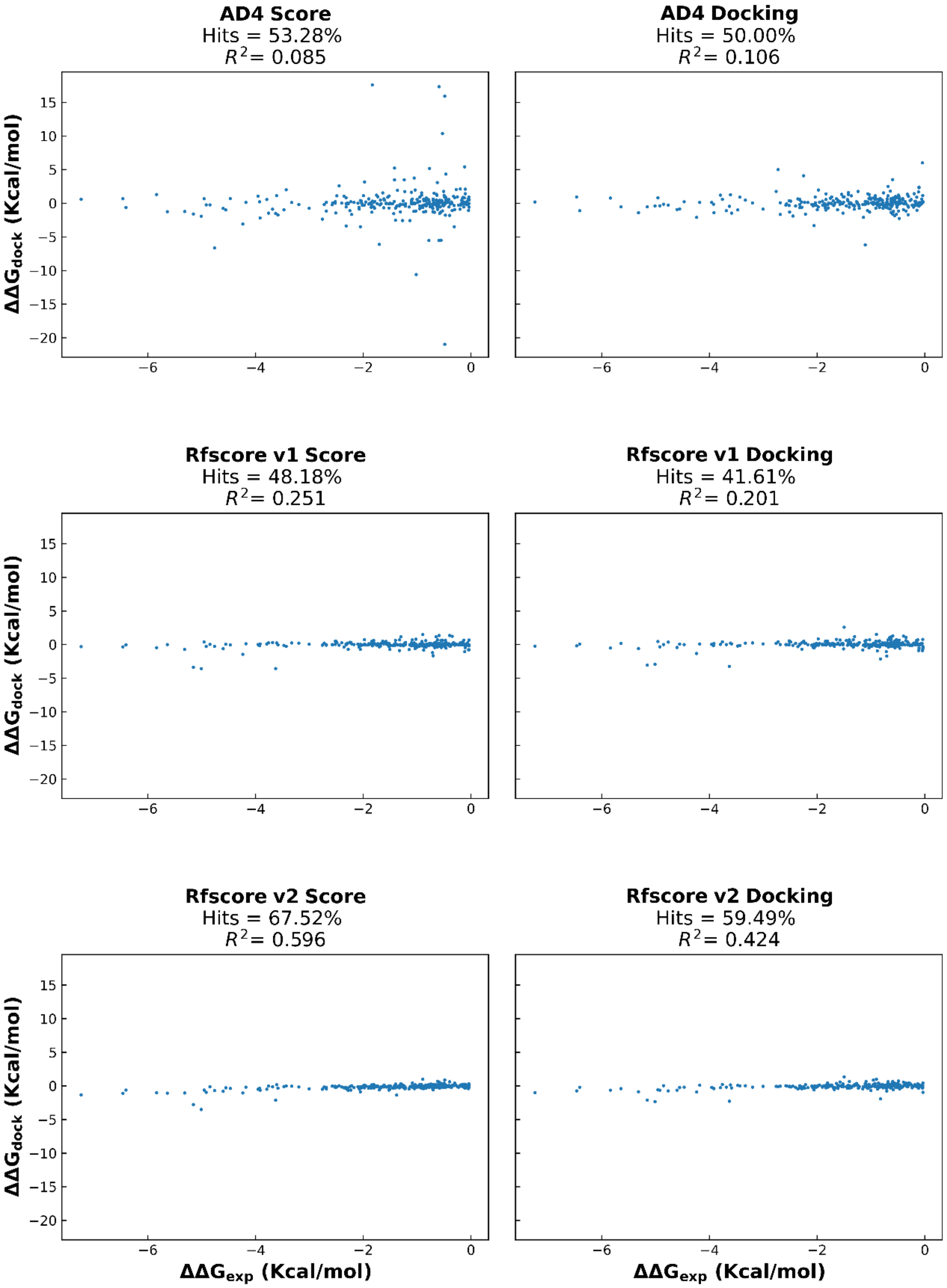


*Figure S2 - Continued (Page 2 of 4).*

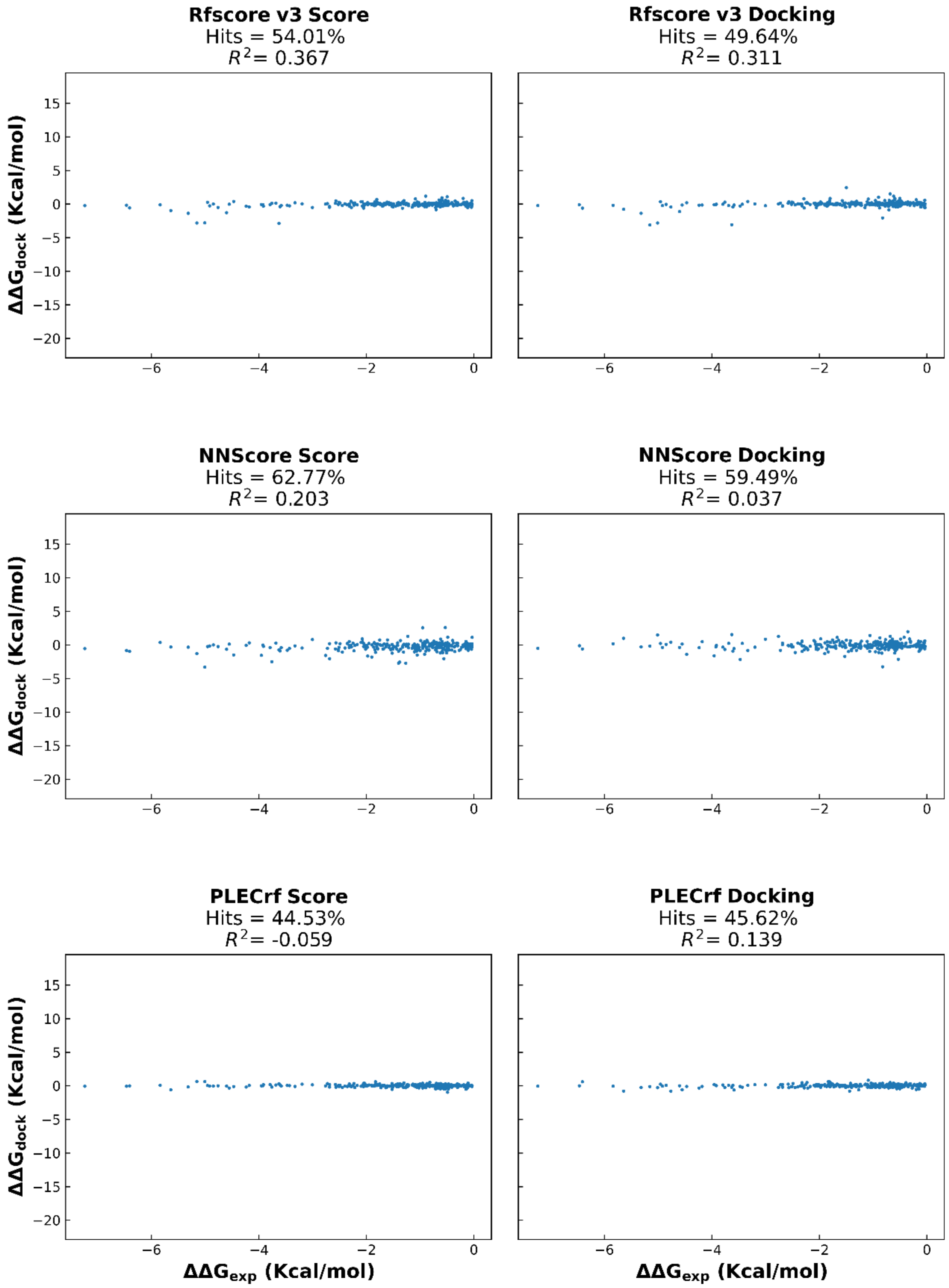


*Figure S2 - Continued (Page 3 of 4).*

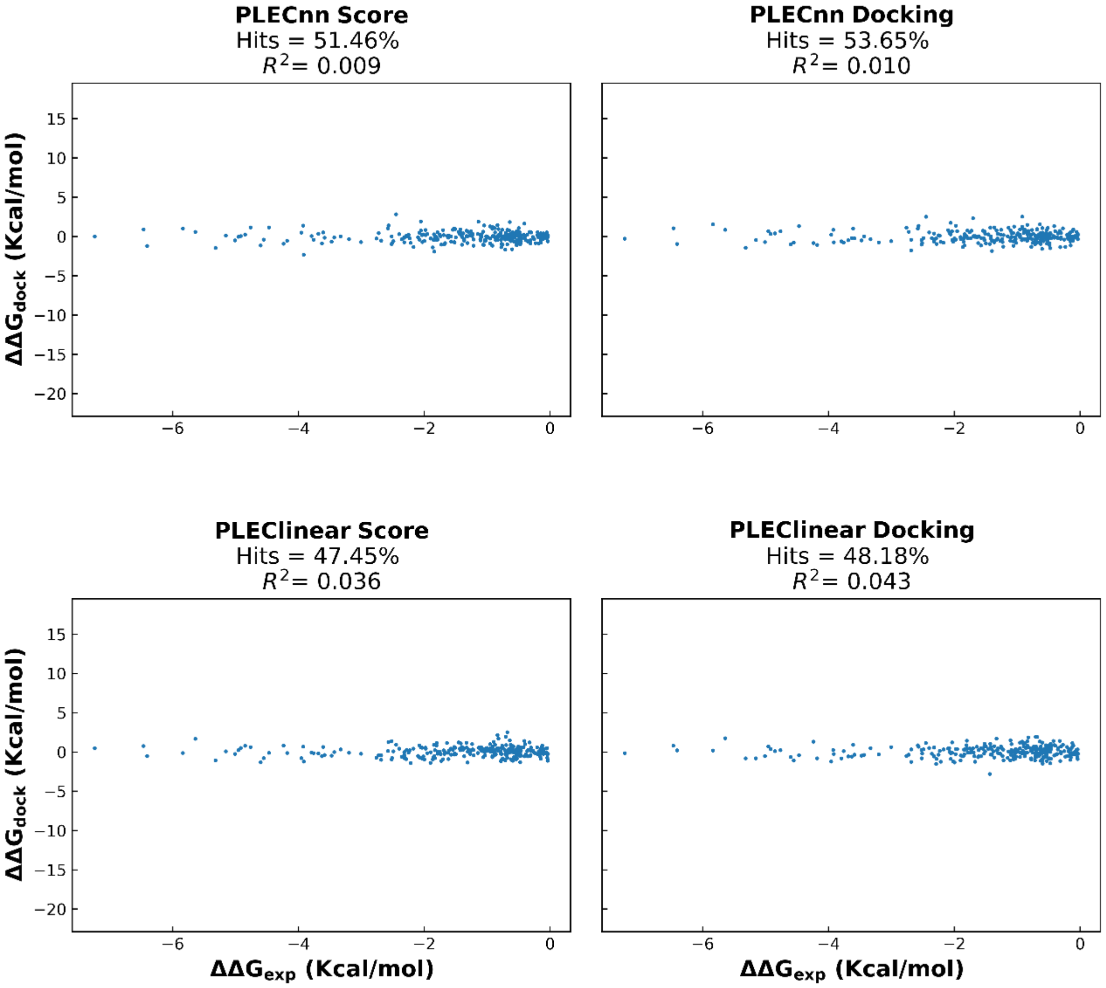


*Figure S2 - Continued (Page 4 of 4).*

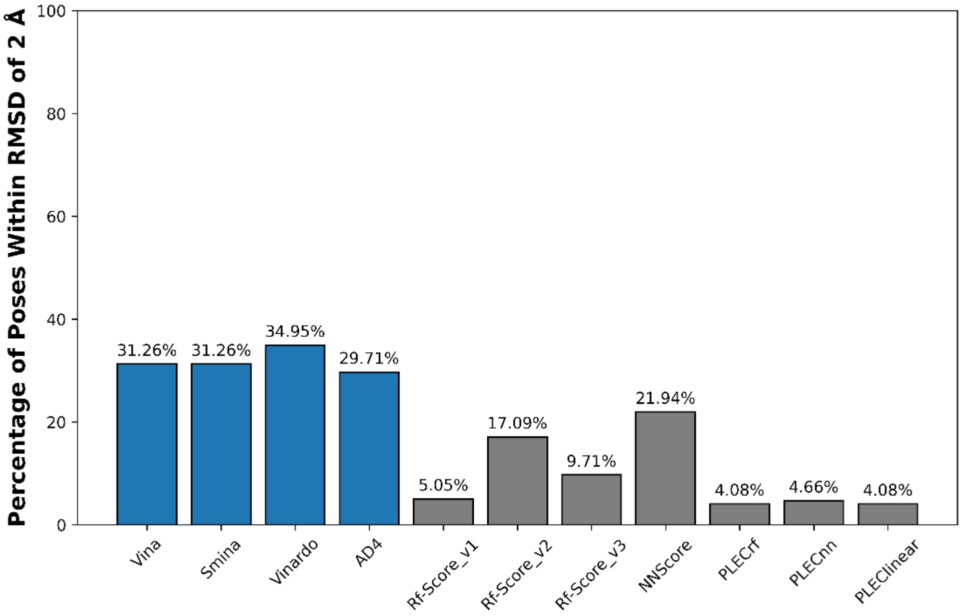


*Figure S3 - Docking power prediction of the results obtained by the platform. Docking ability measured as a percentage of poses, for the 268 pairs of protein-ligand complexes, within a 2 Å RMSD from the crystalized one which the scoring function predicted as best pose. Classical scoring functions are represented in blue and machine learning-based scoring functions in gray.*

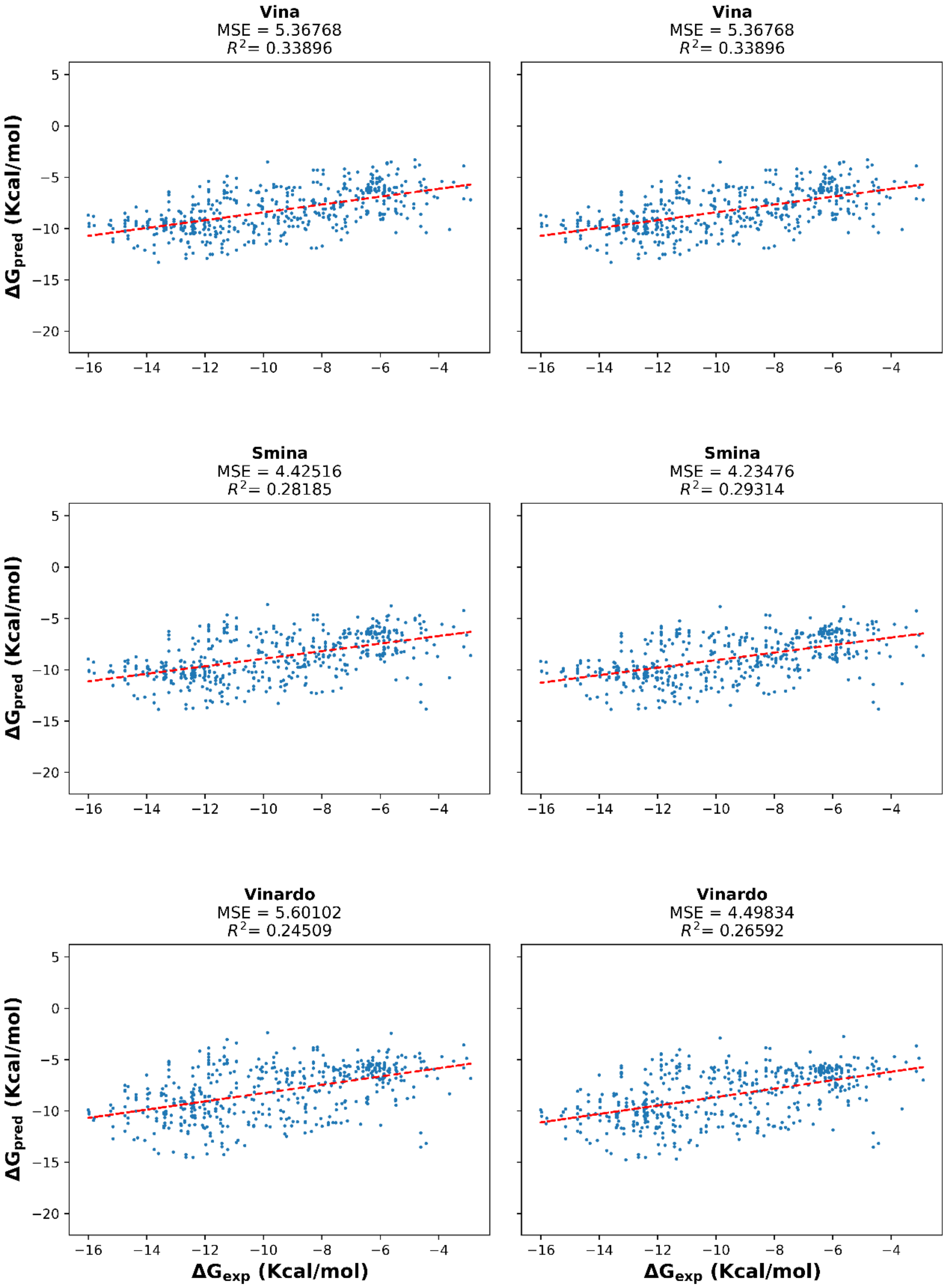


*Figure S4 - Scatter plots comparing predicted binding affinities from the platform's results to experimental binding affinities. Left column - Predicted binding affinities when considering the best pose found by Vina. Right column – predicted binding affinities when considering the best scored pose by each scoring function. The result of a linear regression is represented in red. Structures from PDBbind v2018 database. (Page 1 of 4).*

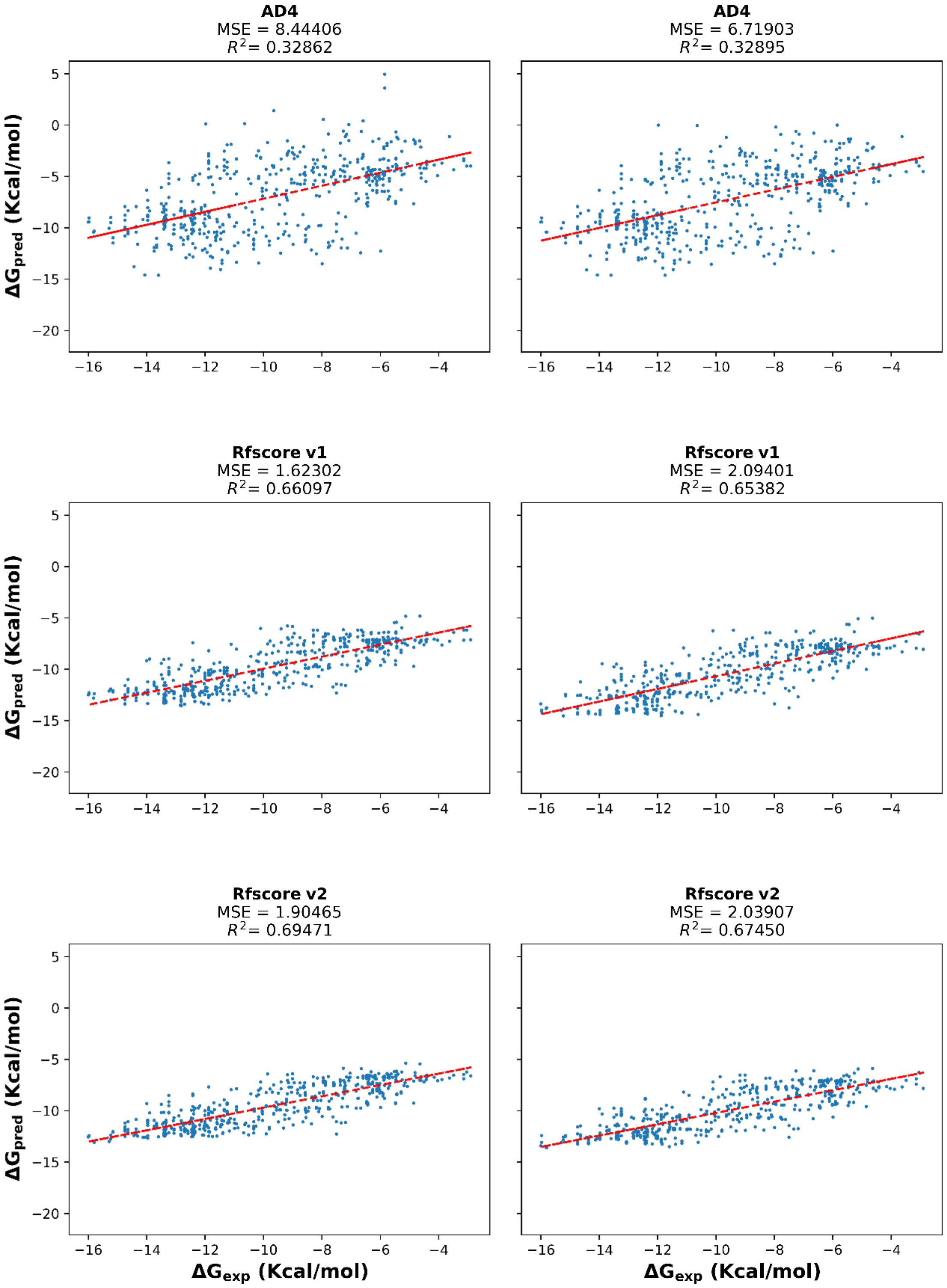


*Figure S4 - Continued. (Page 2 of 4).*

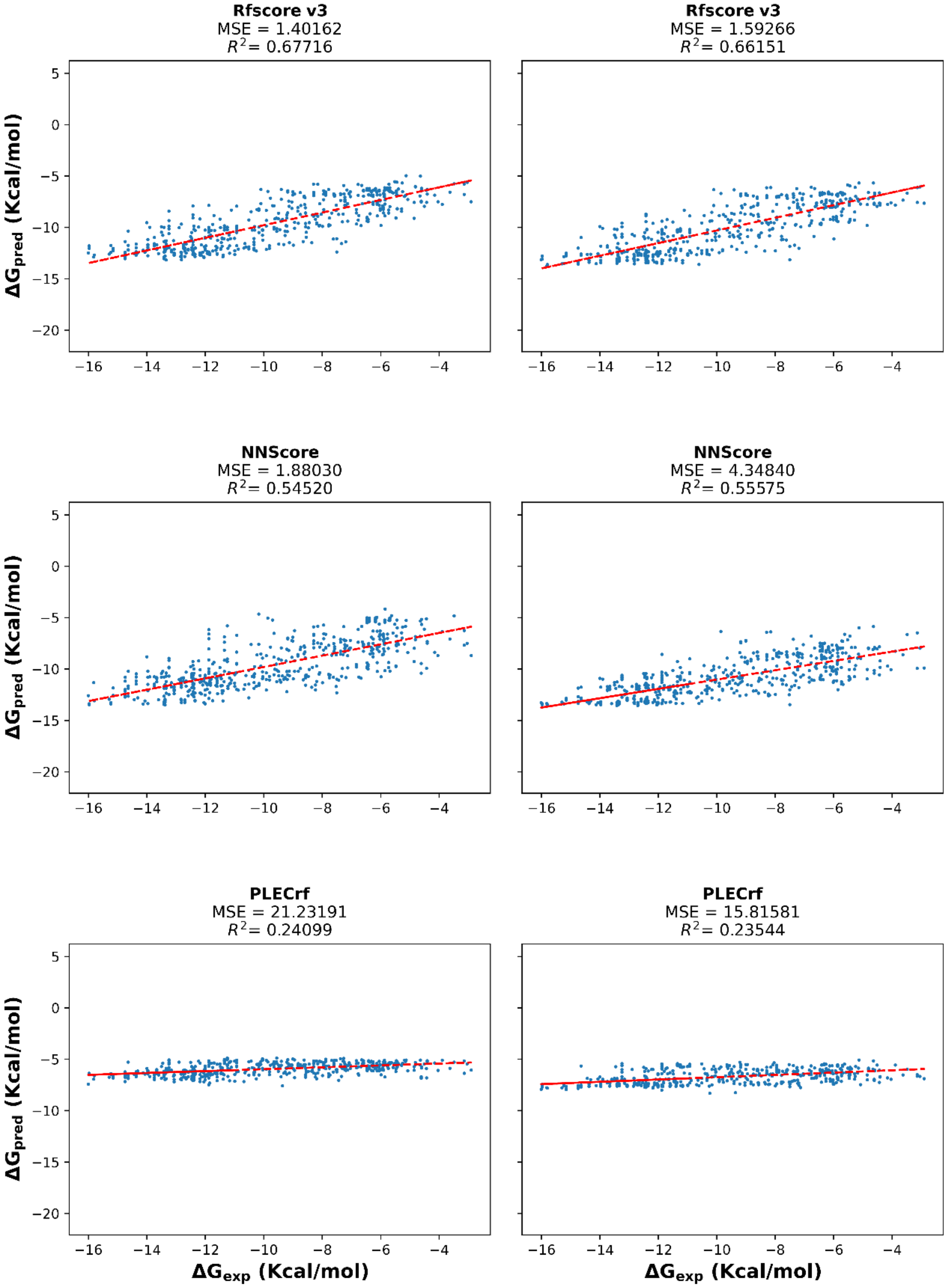


*Figure S4 - Continued. (Page 3 of 4).*

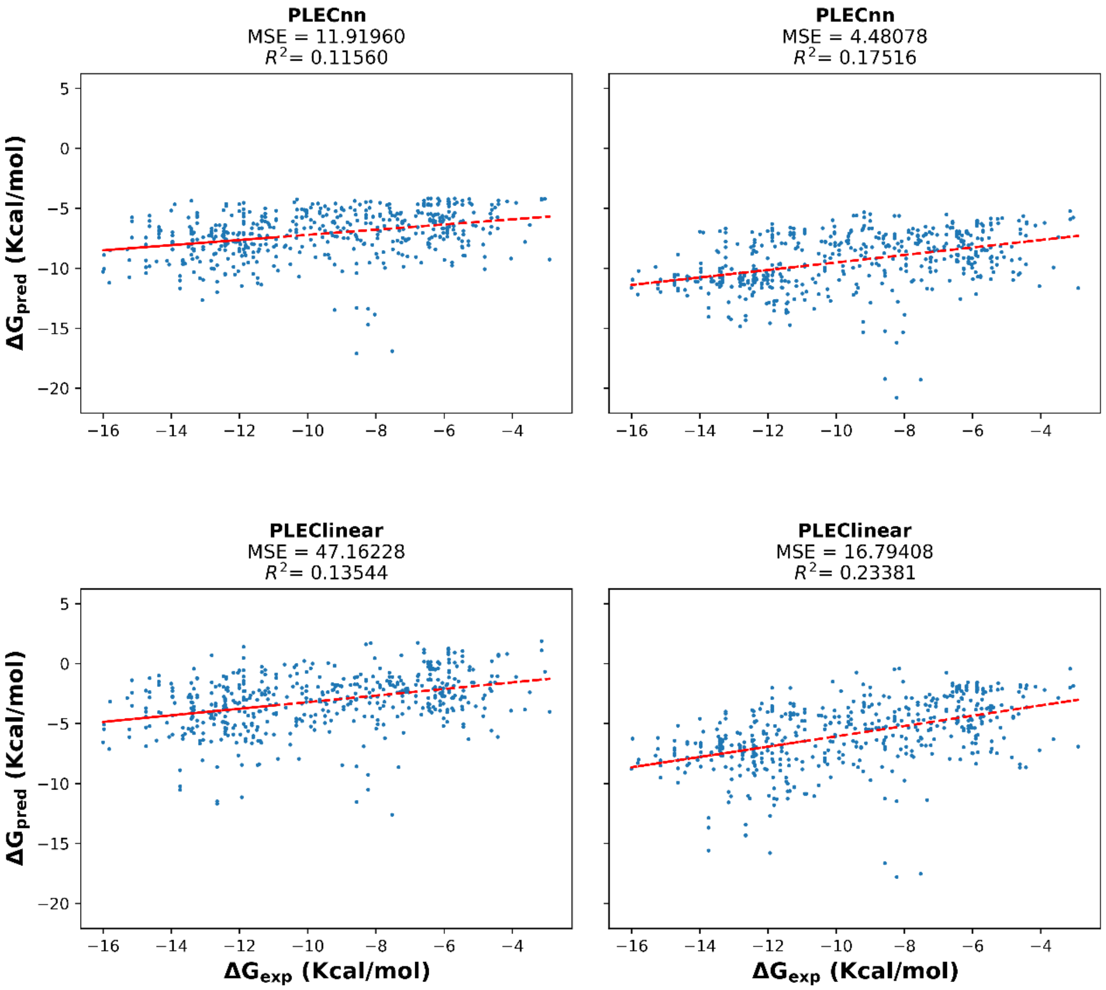


*Figure S4 - Continued. (Page 4 of 4).*

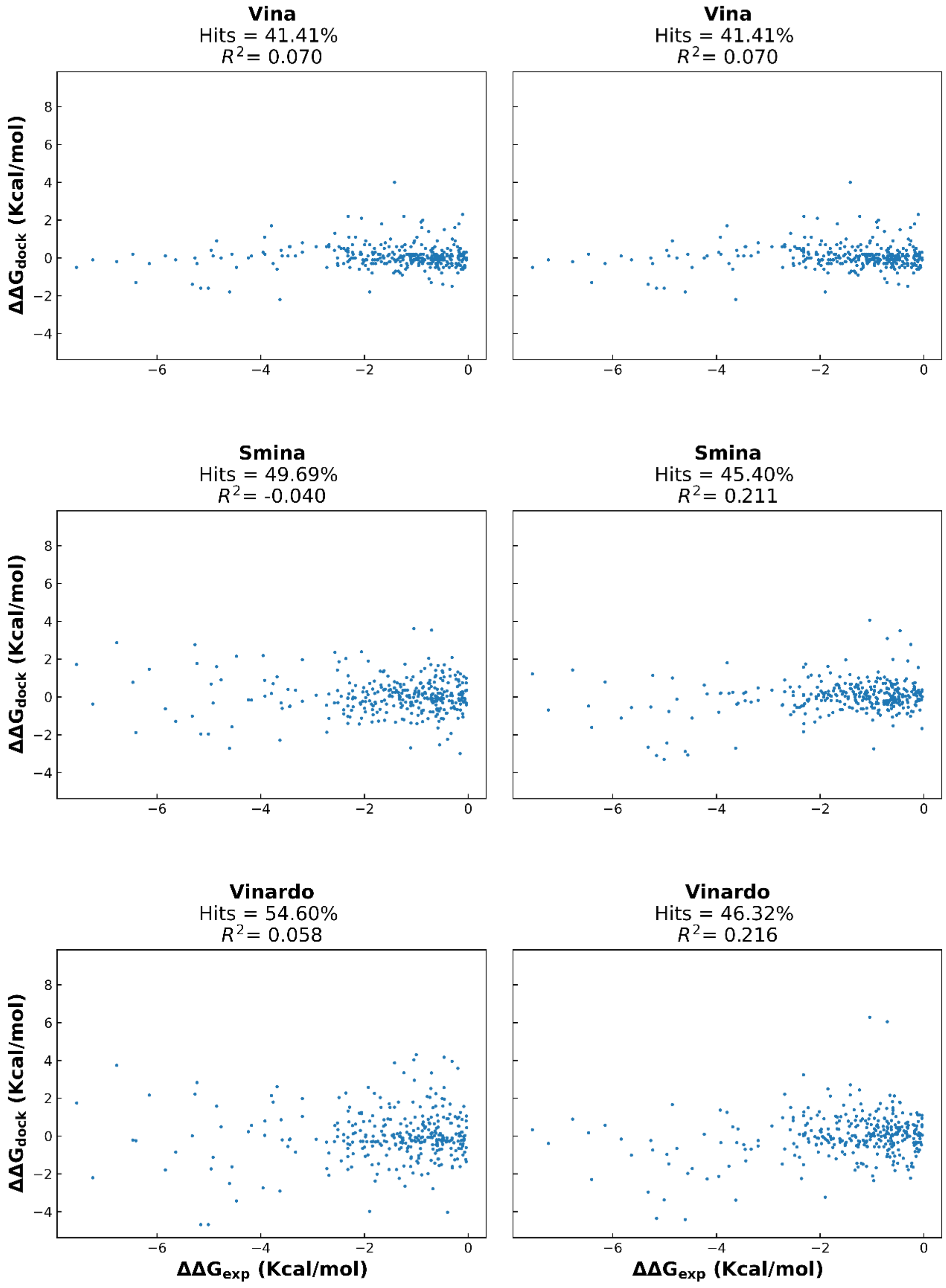


*Figure S5 - Scatter plots comparing predicted ΔΔG from the platform's results to experimental ΔΔG. Left column - Predicted ΔΔG when considering the best pose found by Vina. Right column – predicted ΔΔG when considering the best scored pose by each scoring function. Hits – Percentage of successes on predicting the mutation's effect on binding. Structures from PDBbind v2018 database. (Page 1 of 4).*

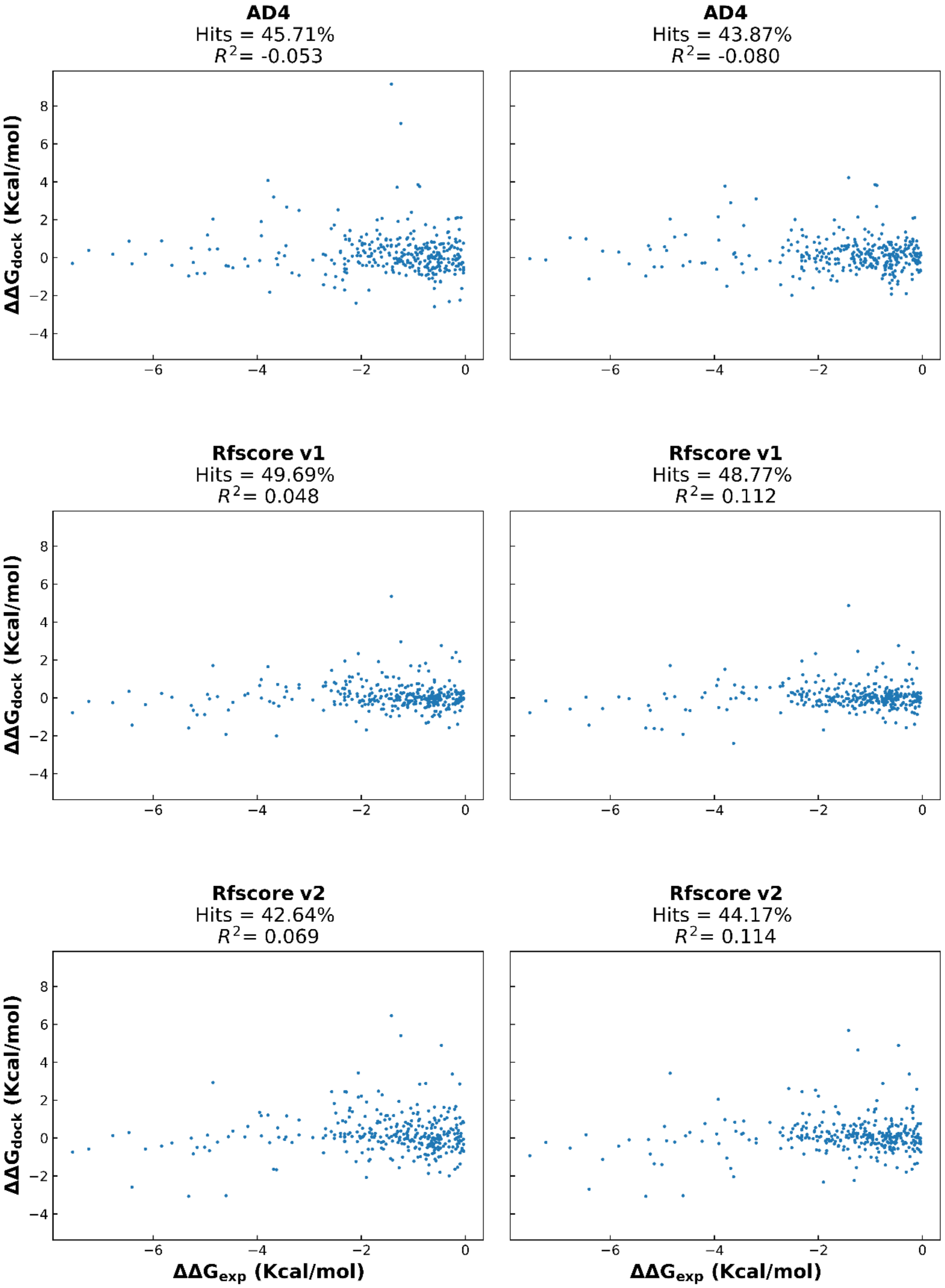


*Figure S5 - Continued (Page 2 of 4).*

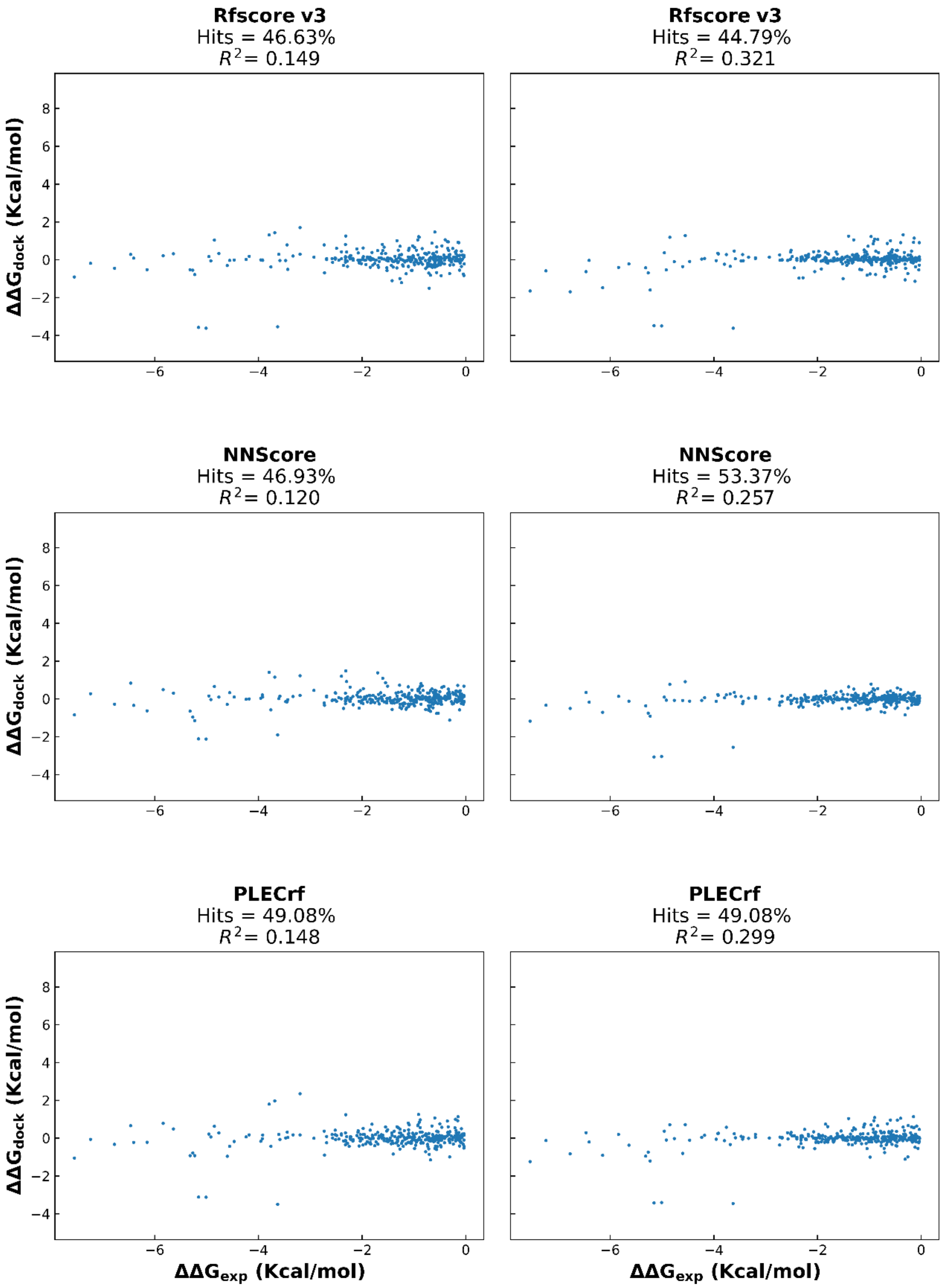


*Figure S5 - Continued (Page 3 of 4).*

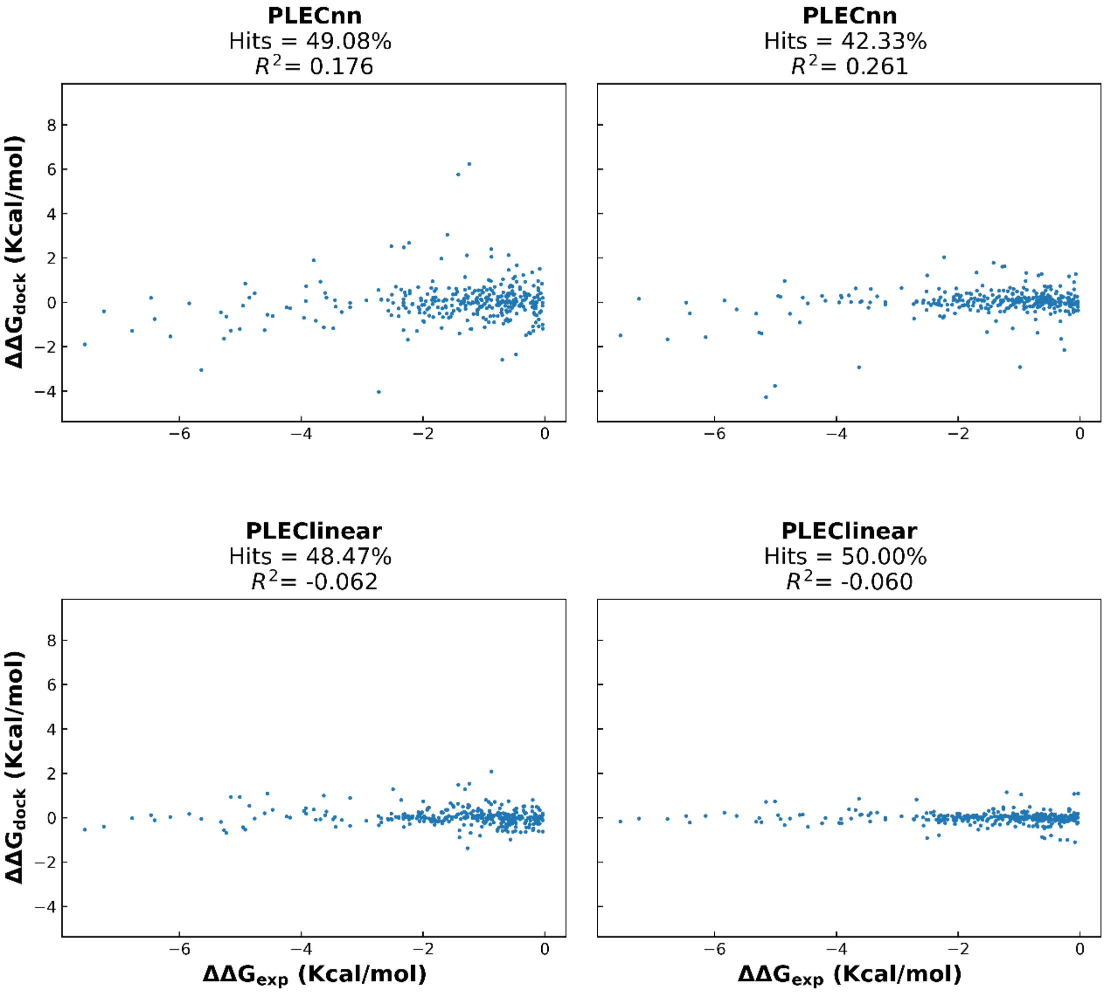


*Figure S5 - Continued (Page 4 of 4).*

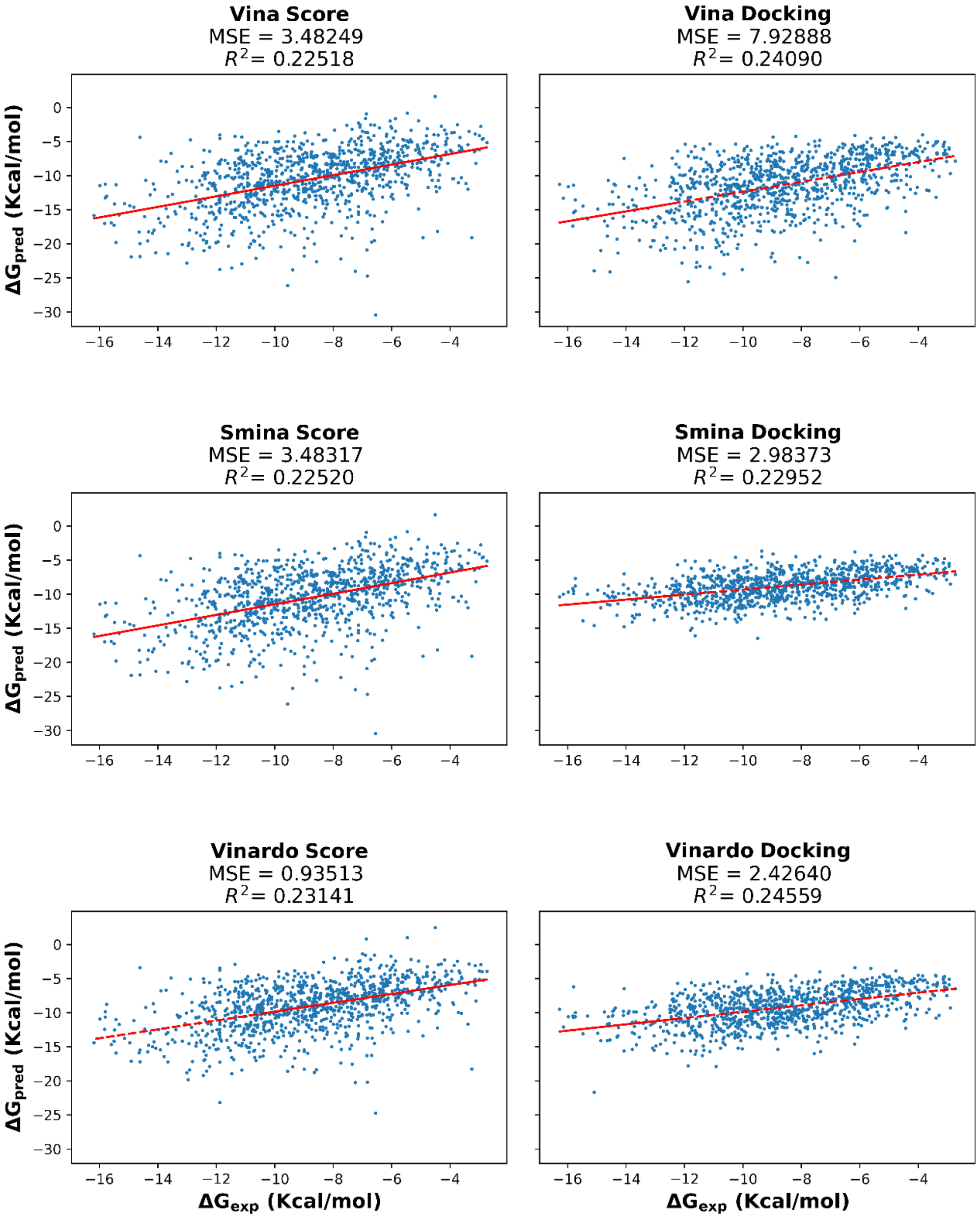


*Figure S6 - Scatter plots comparing predicted binding affinities to experimental binding affinities for the stacking validation set. Left column - Predicted binding affinities when calculated using each scoring function on the crystallized pose (first phase). Right column – predicted binding affinities after re-docking the ligand in the protein's binding pocket (second phase). The result of a linear regression is represented in red. Structures from PDBbind v2018 database. (Page 1 of 4).*

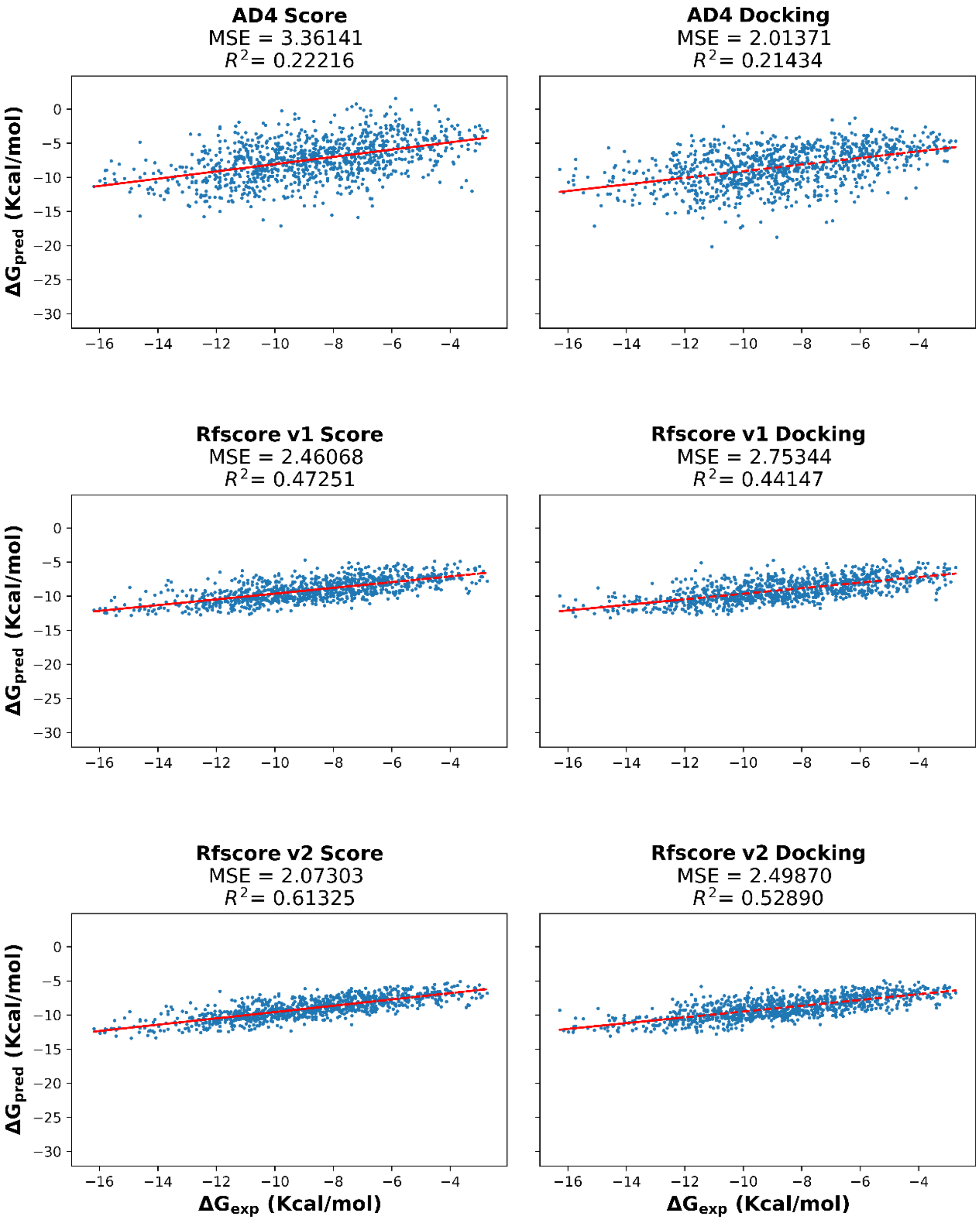


*Figure S6 - Continued. (Page 2 of 4).*

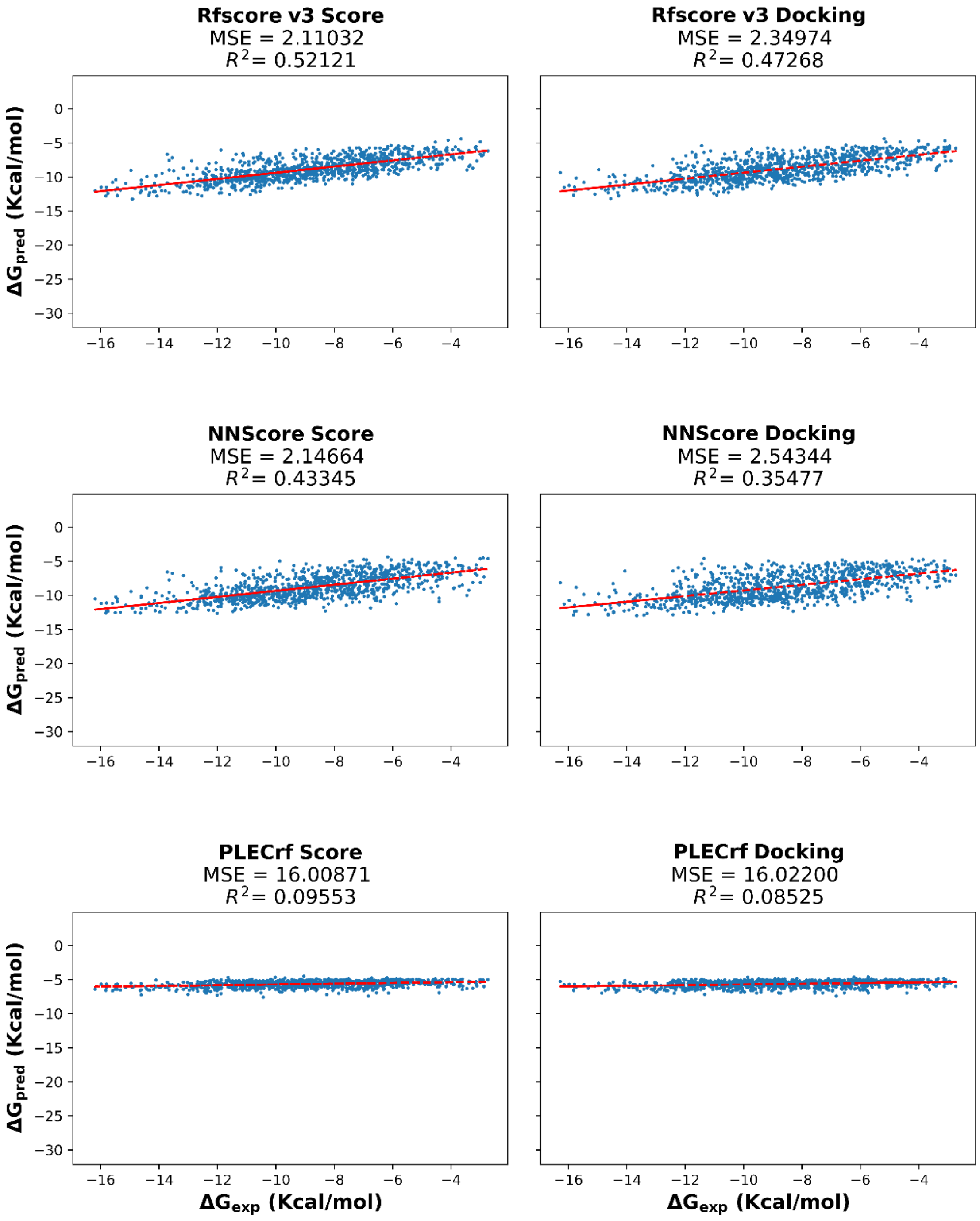


*Figure S6 - Continued. (Page 3 of 4).*

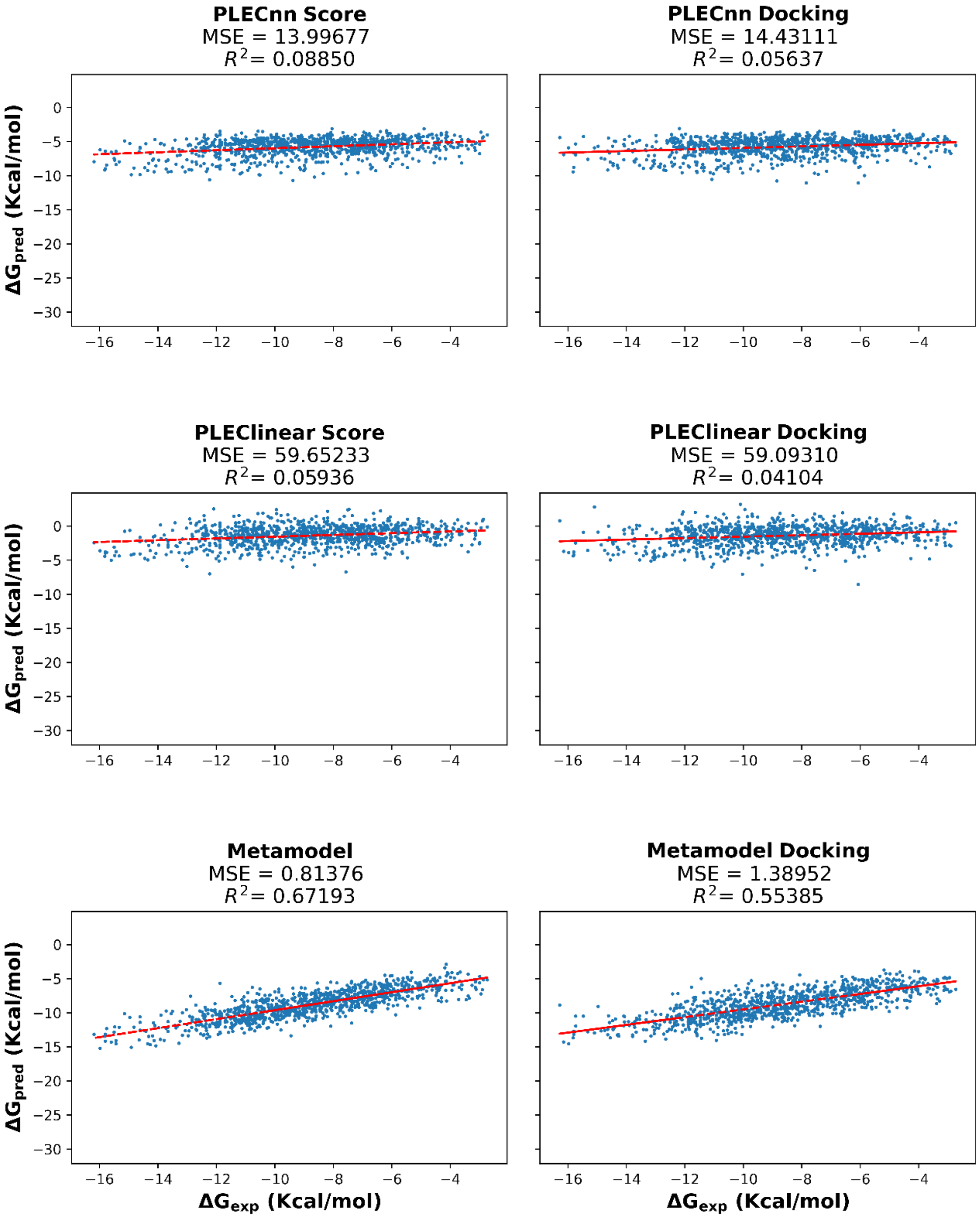


*Figure S6 - Continued. (Page 4 of 4).*

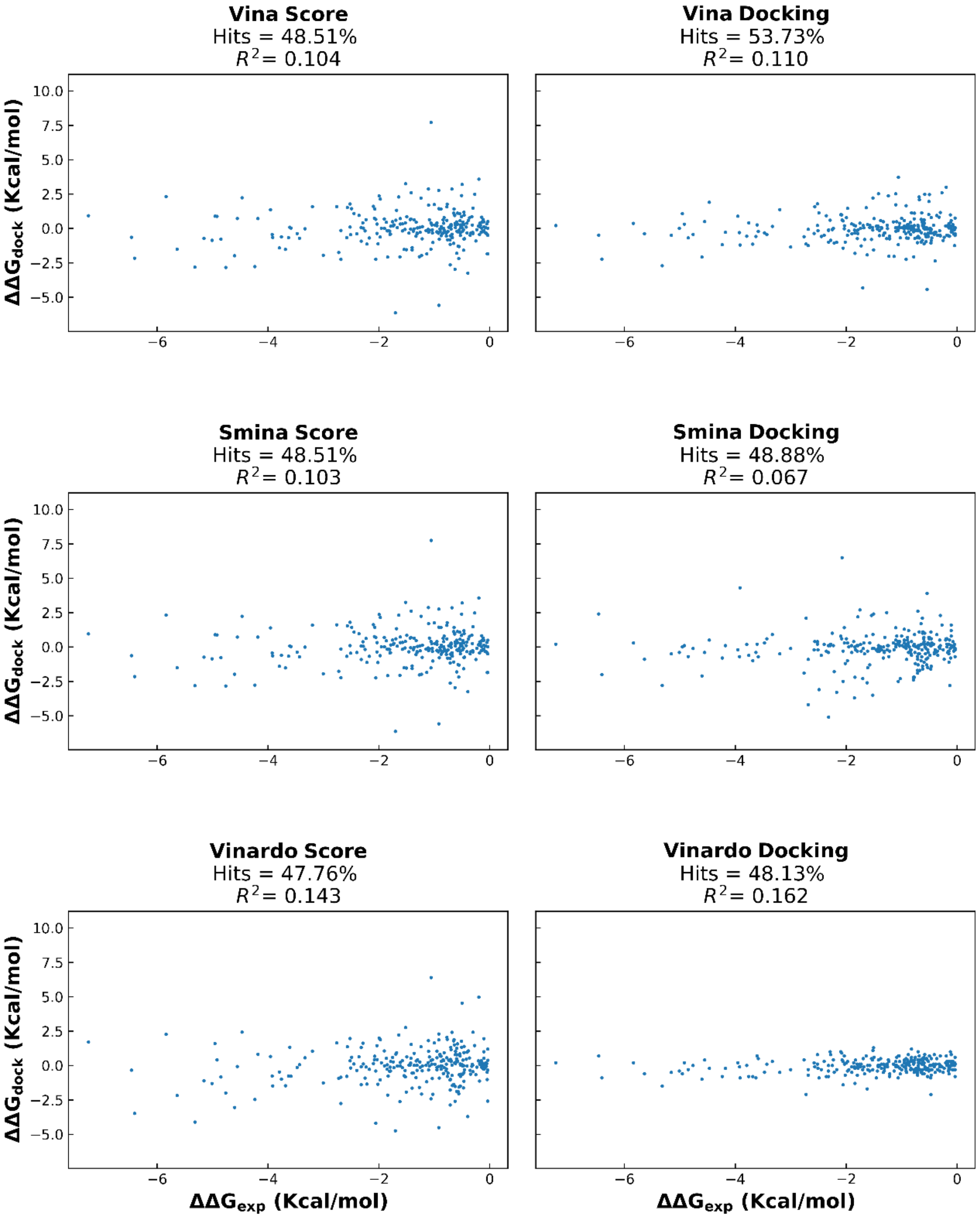


*Figure S7 - Scatter plots comparing predicted ΔΔG to experimental ΔΔG for the stacking validation set. Left column - Predicted ΔΔG when calculated using each scoring function on the crystallized pose (first phase). Right column – predicted ΔΔG after re-docking the ligand in the protein's binding pocket (second phase). Hits – Percentage of successes on predicting the mutation's effect on binding. Structures from PDBbind v2018 database. (Page 1 of 4).*

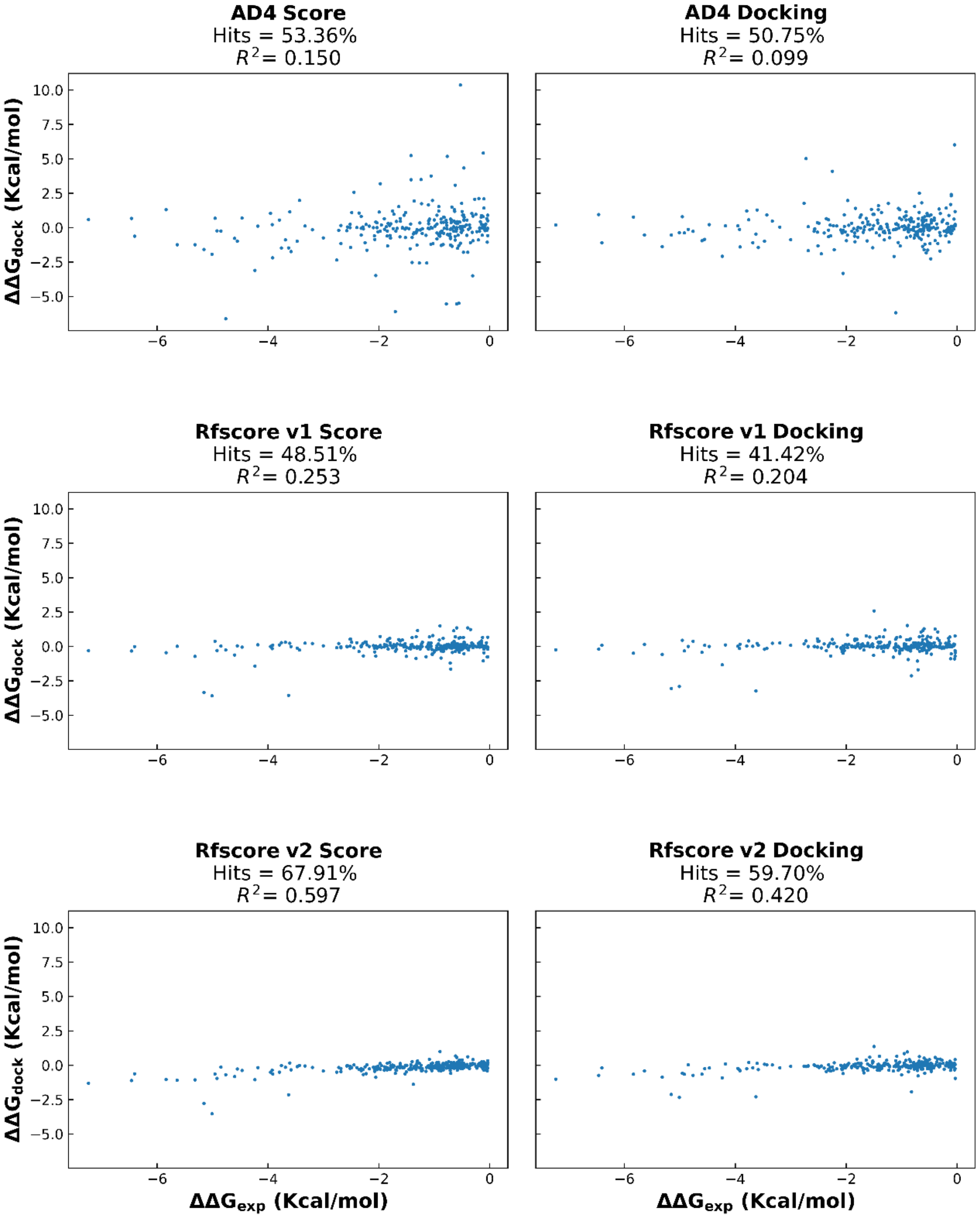


*Figure S7 - Continued (Page 2 of 4).*

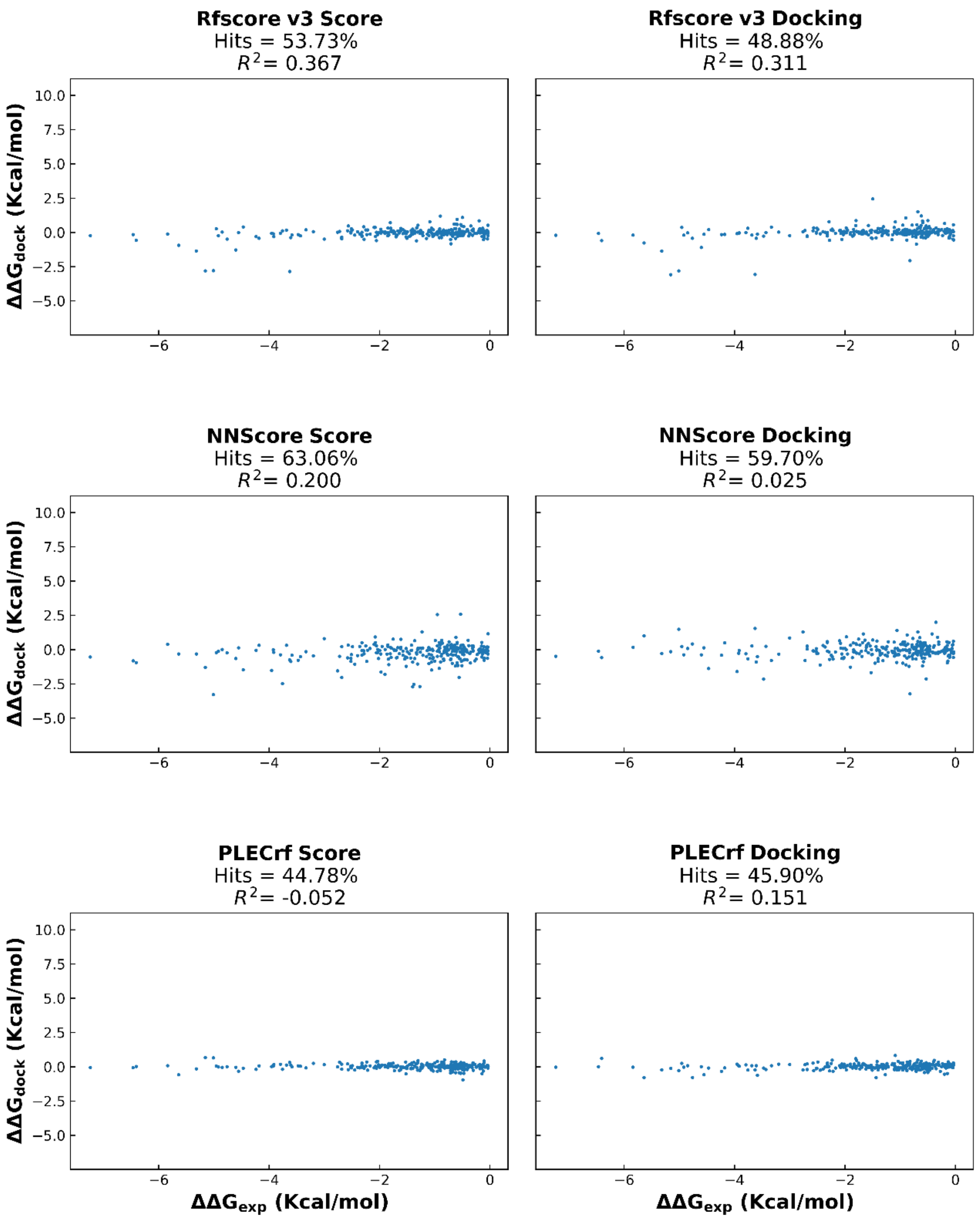


*Figure S7 - Continued (Page 3 of 4).*

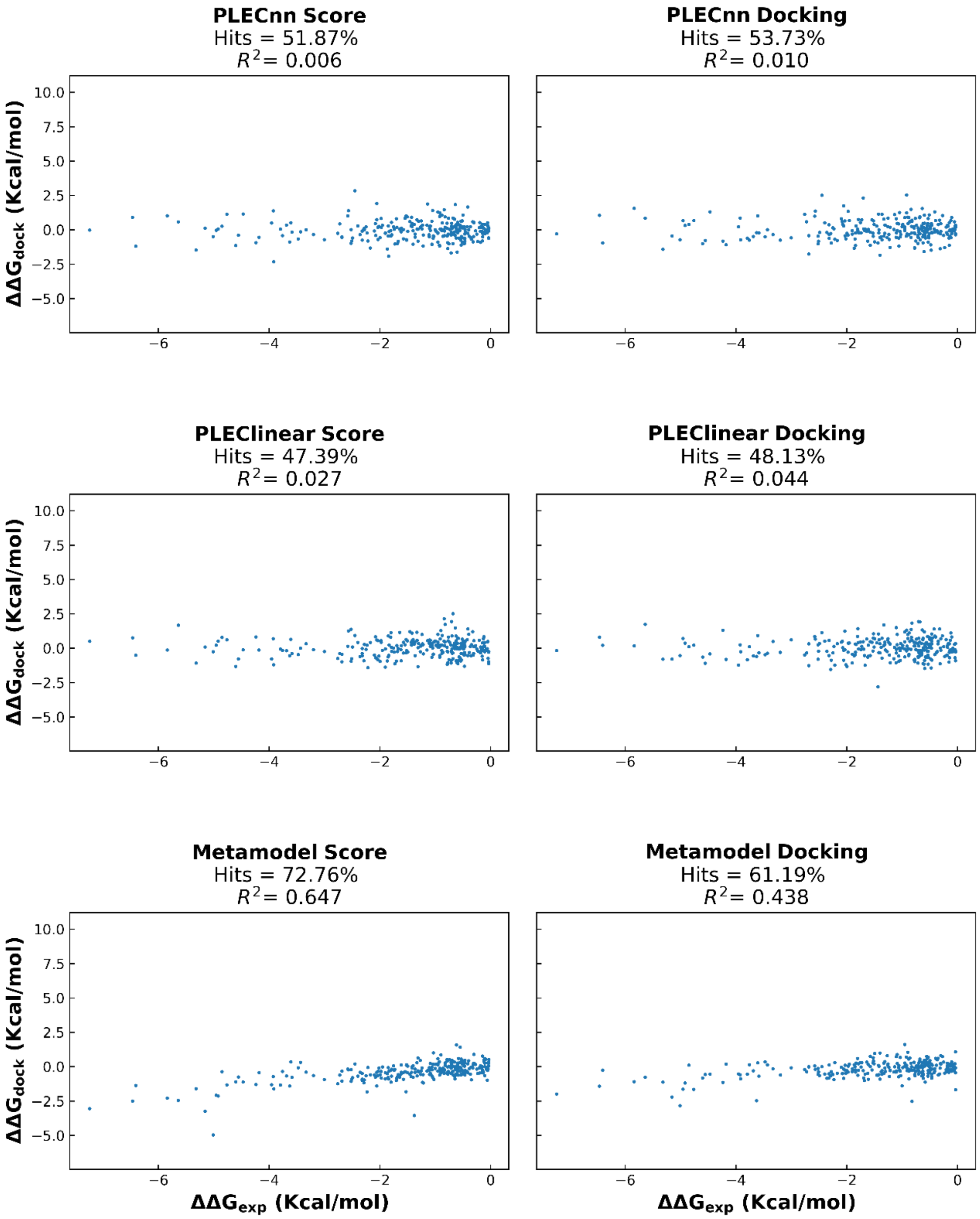


*Figure S7 - Continued (Page 4 of 4).*

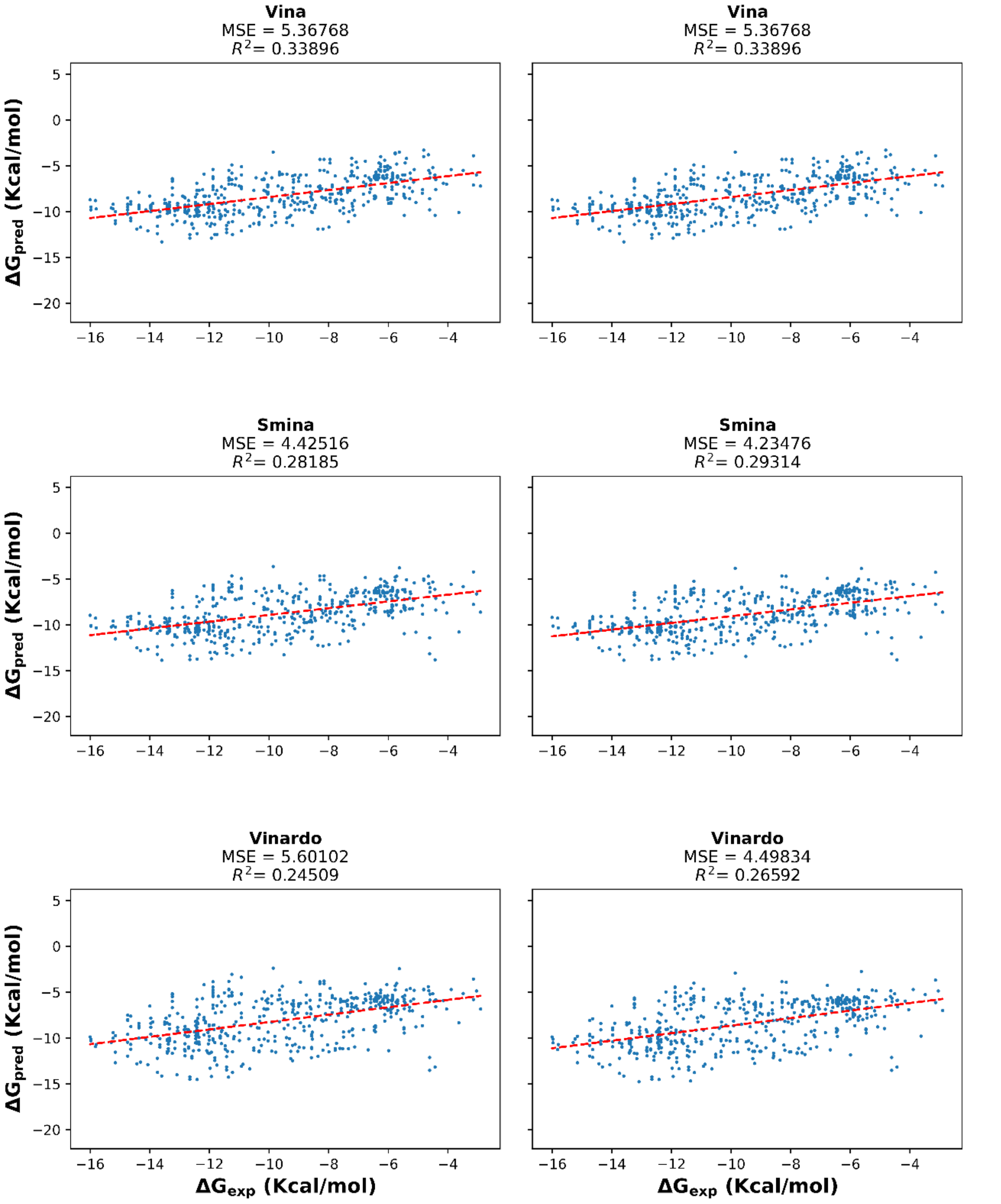


*Figure S5 - Scatter plots comparing predicted binding affinities from the platform's results to experimental binding affinities for the stacking validation set. Left column - Predicted binding affinities when considering the best pose found by Vina. Right column – predicted binding affinities when considering the best scored pose by each scoring function. The result of a linear regression is represented in red. Structures from PDBbind v2018 database. (Page 1 of 4).*

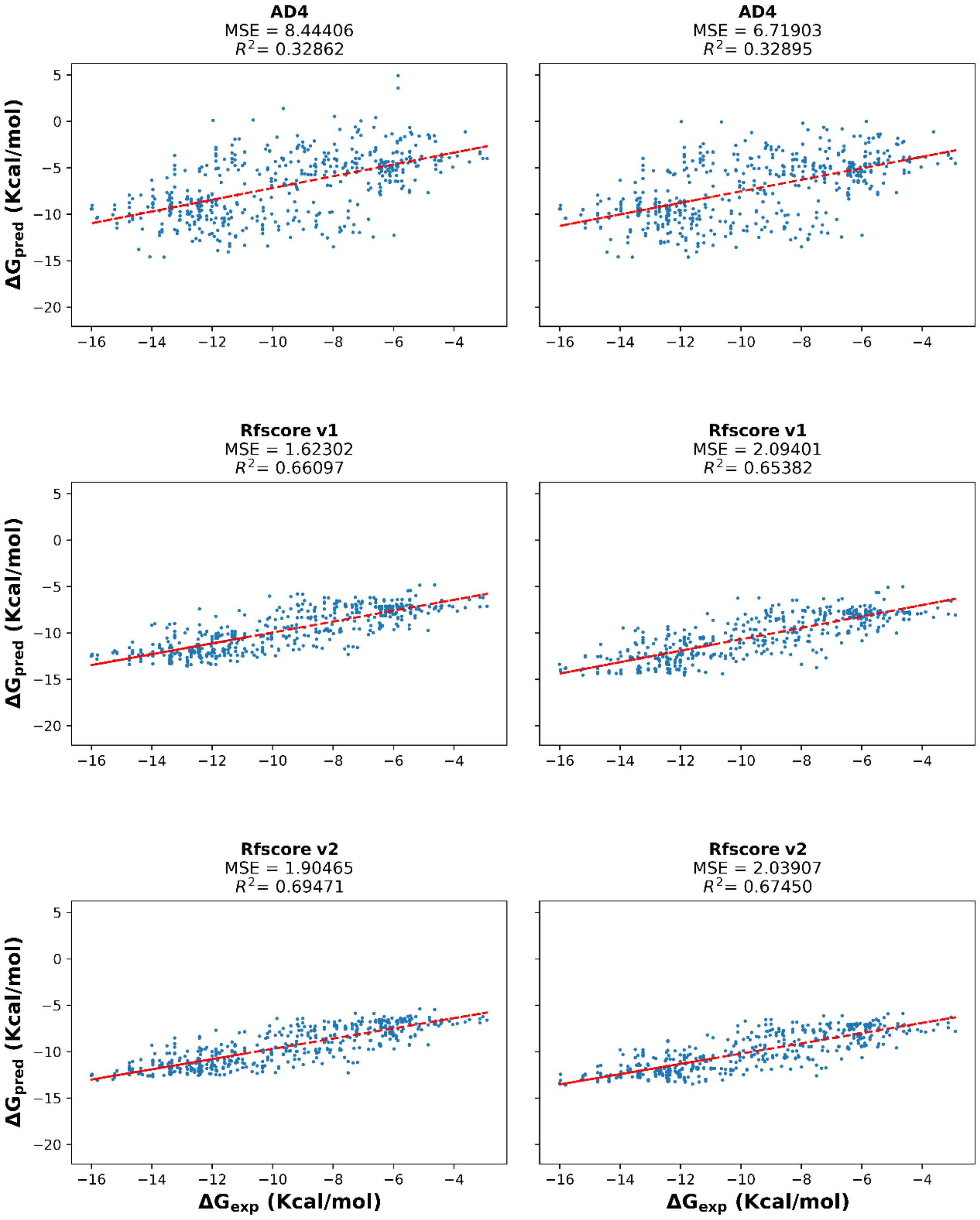


*Figure S8 - Continued. (Page 2 of 4).*

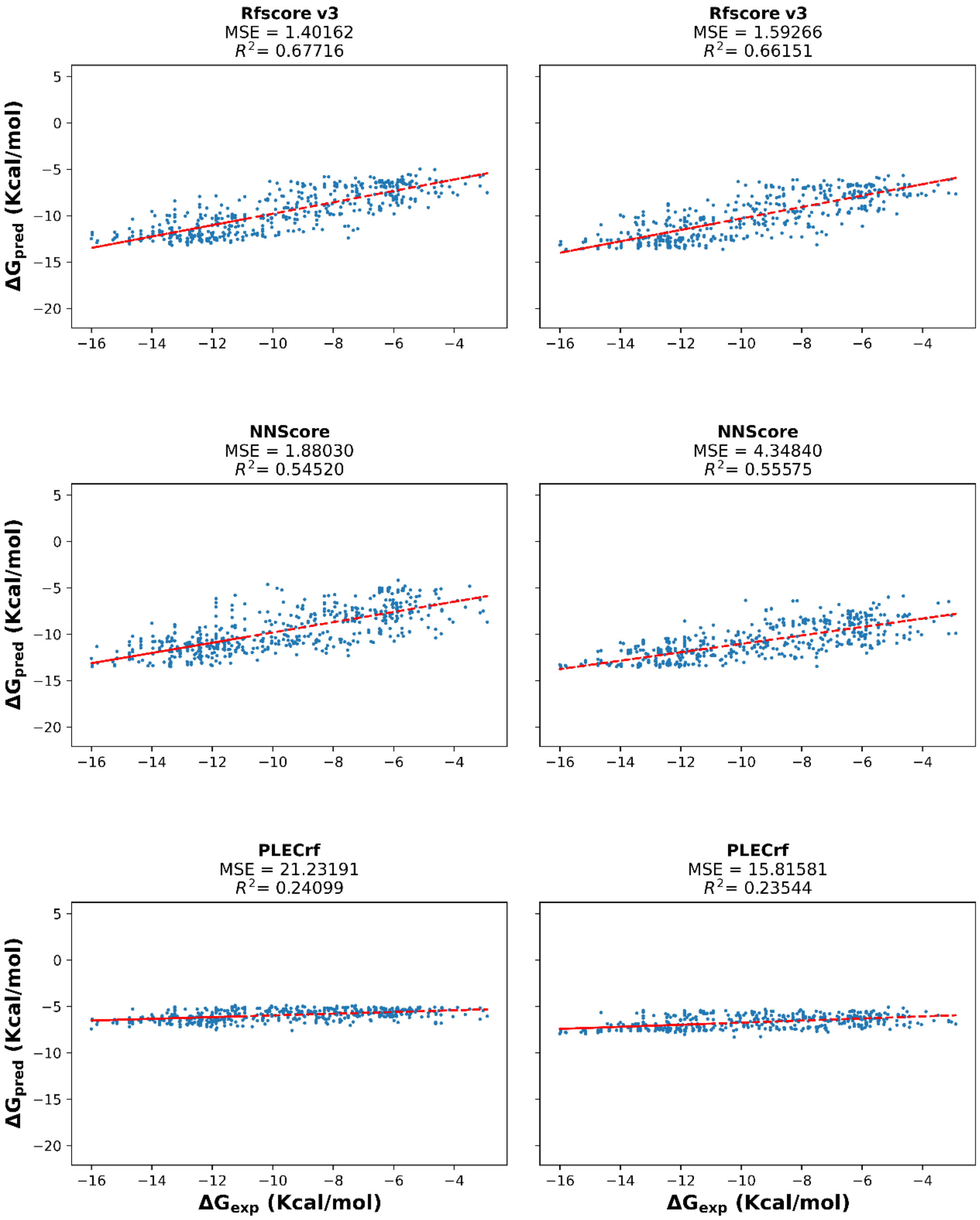


*Figure S8 - Continued. (Page 3 of 4).*

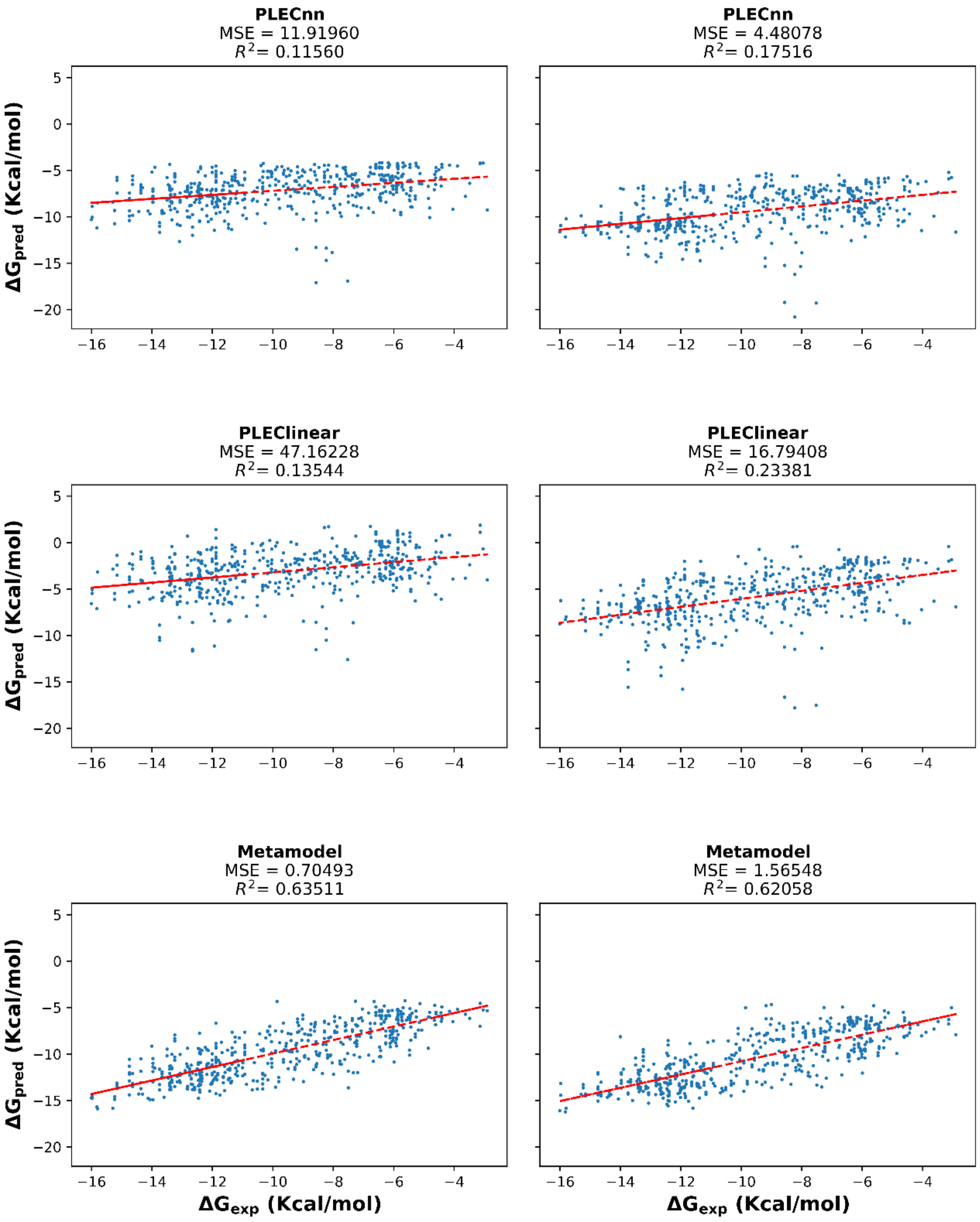


*Figure S8 - Continued. (Page 4 of 4).*

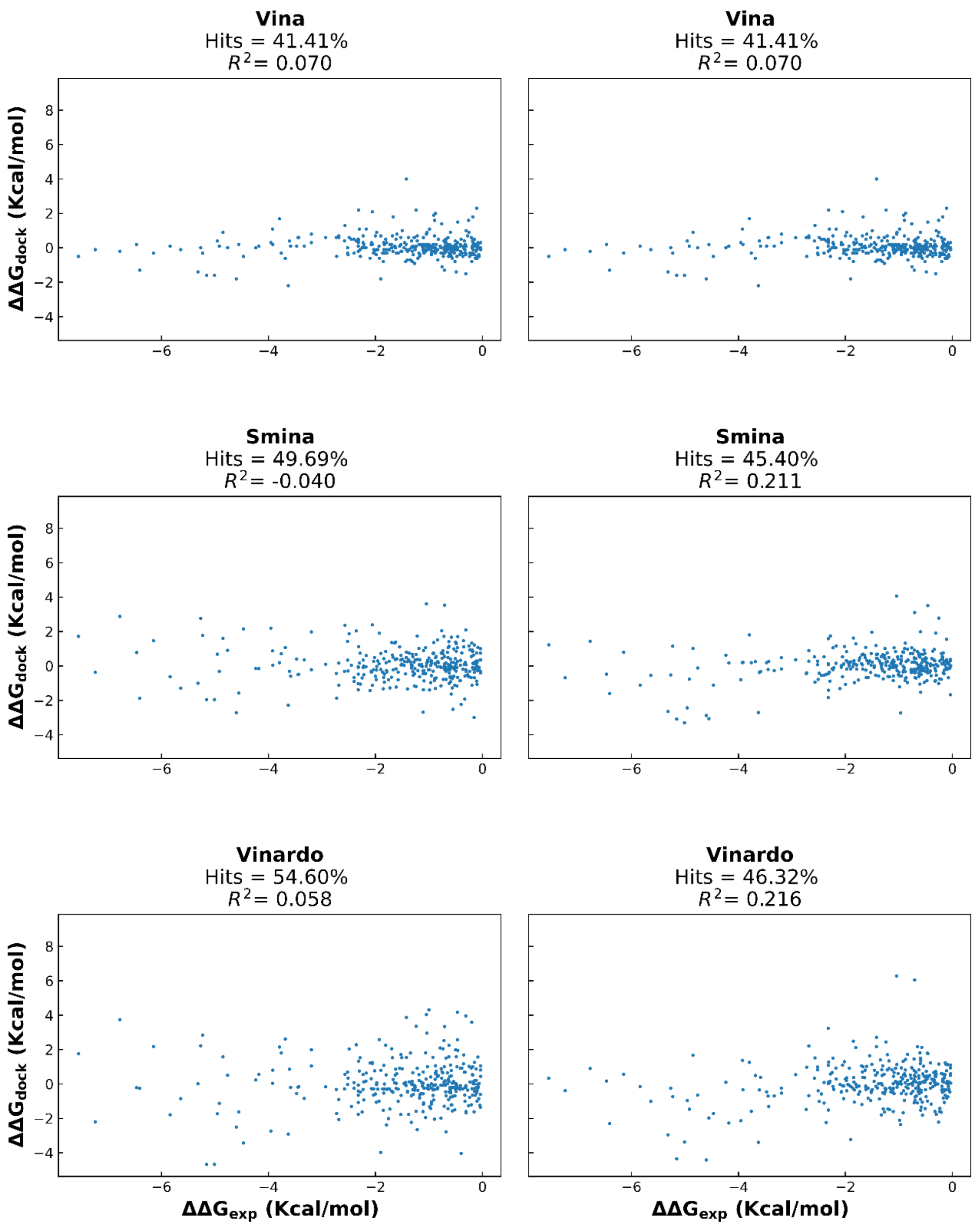


*Figure S9 - Scatter plots comparing predicted ΔΔG from the platform's results to experimental ΔΔG for the stacking validation set. Left column - Predicted ΔΔG when considering the best pose found by Vina. Right column – predicted ΔΔG when considering the best scored pose by each scoring function. Hits – Percentage of successes on predicting the mutation's effect on binding. Structures from PDBbind v2018 database. (Page 1 of 4).*

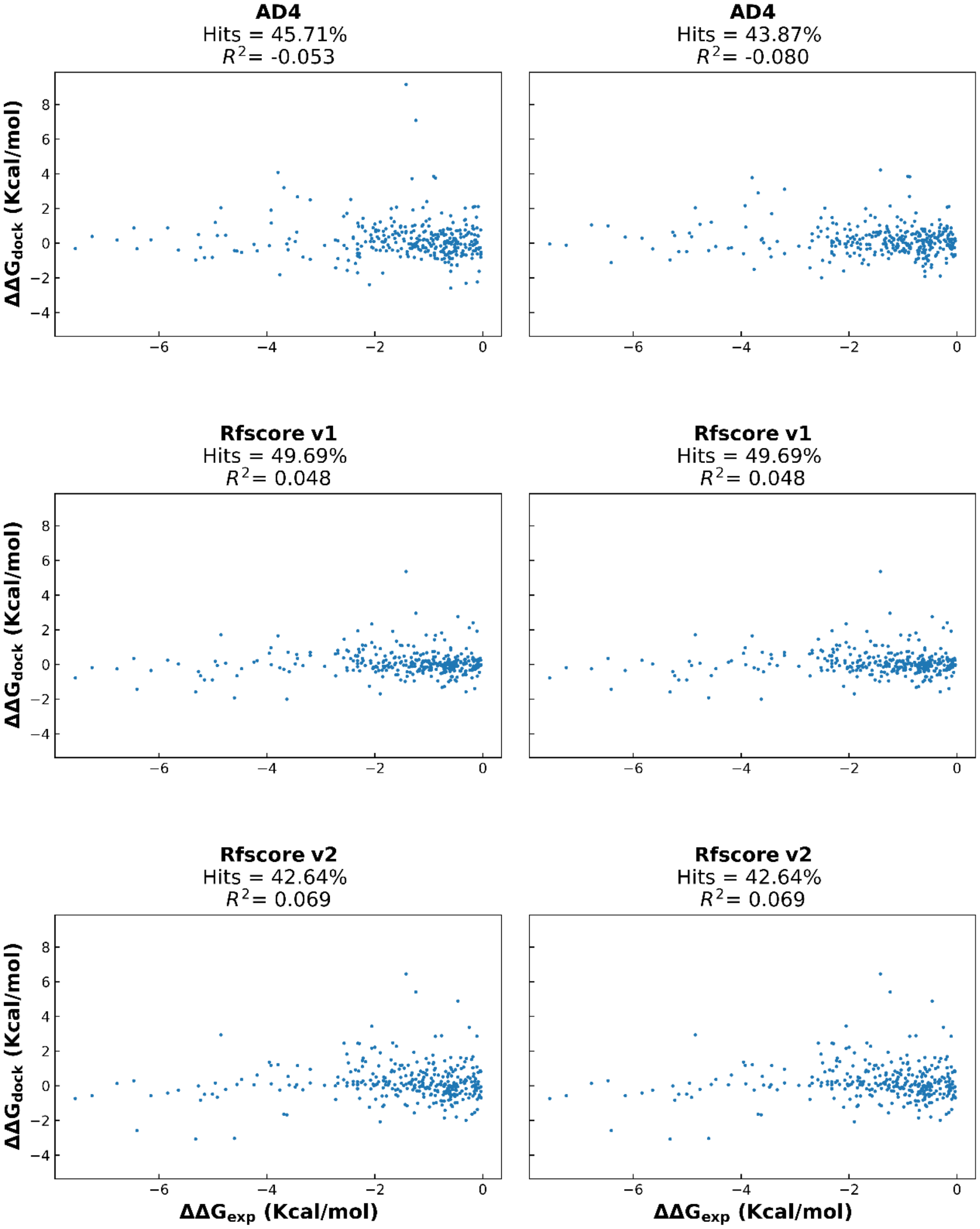


*Figure S9 - Continued. (Page 2 of 4).*

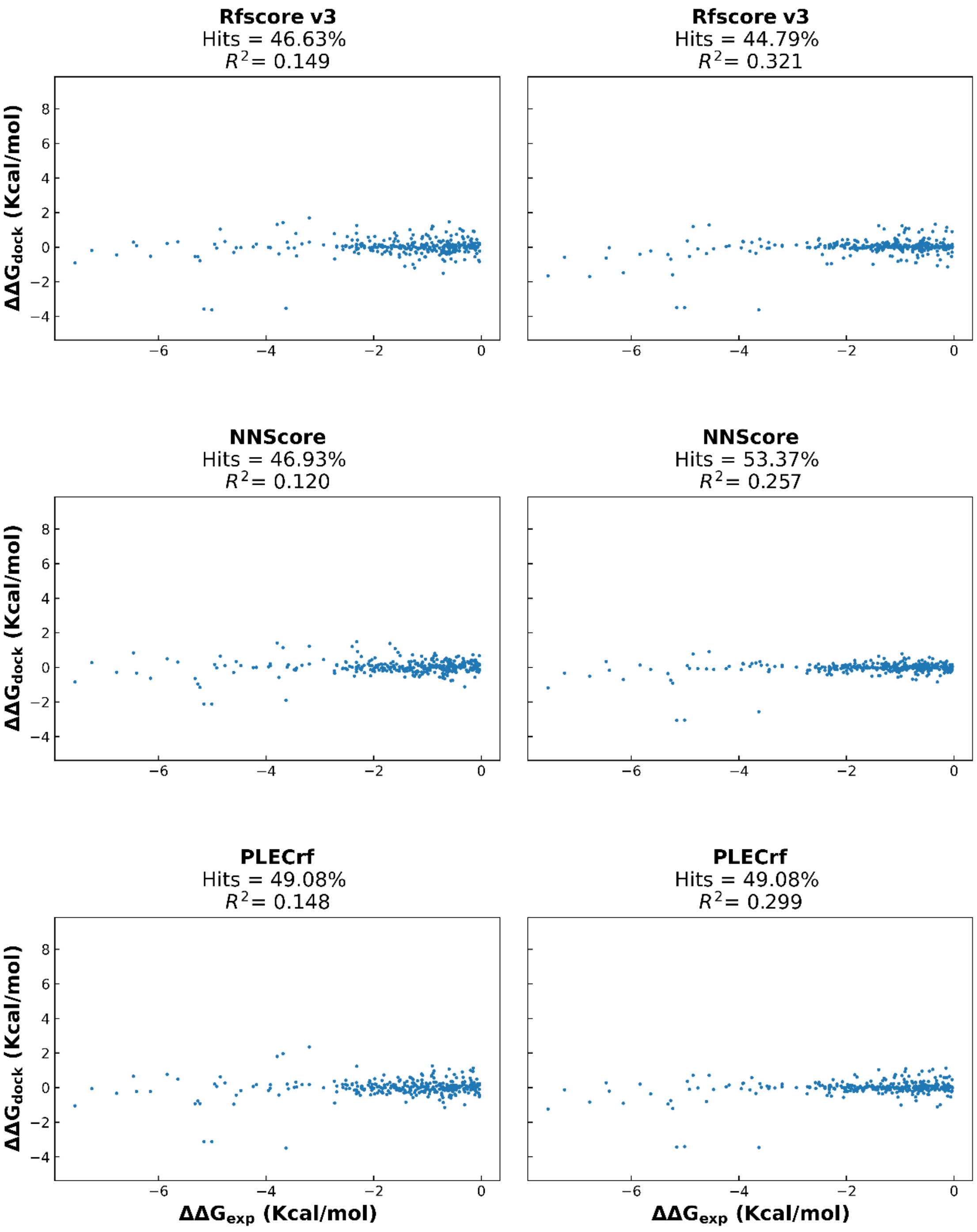


*Figure S9 - Continued. (Page 3 of 4).*

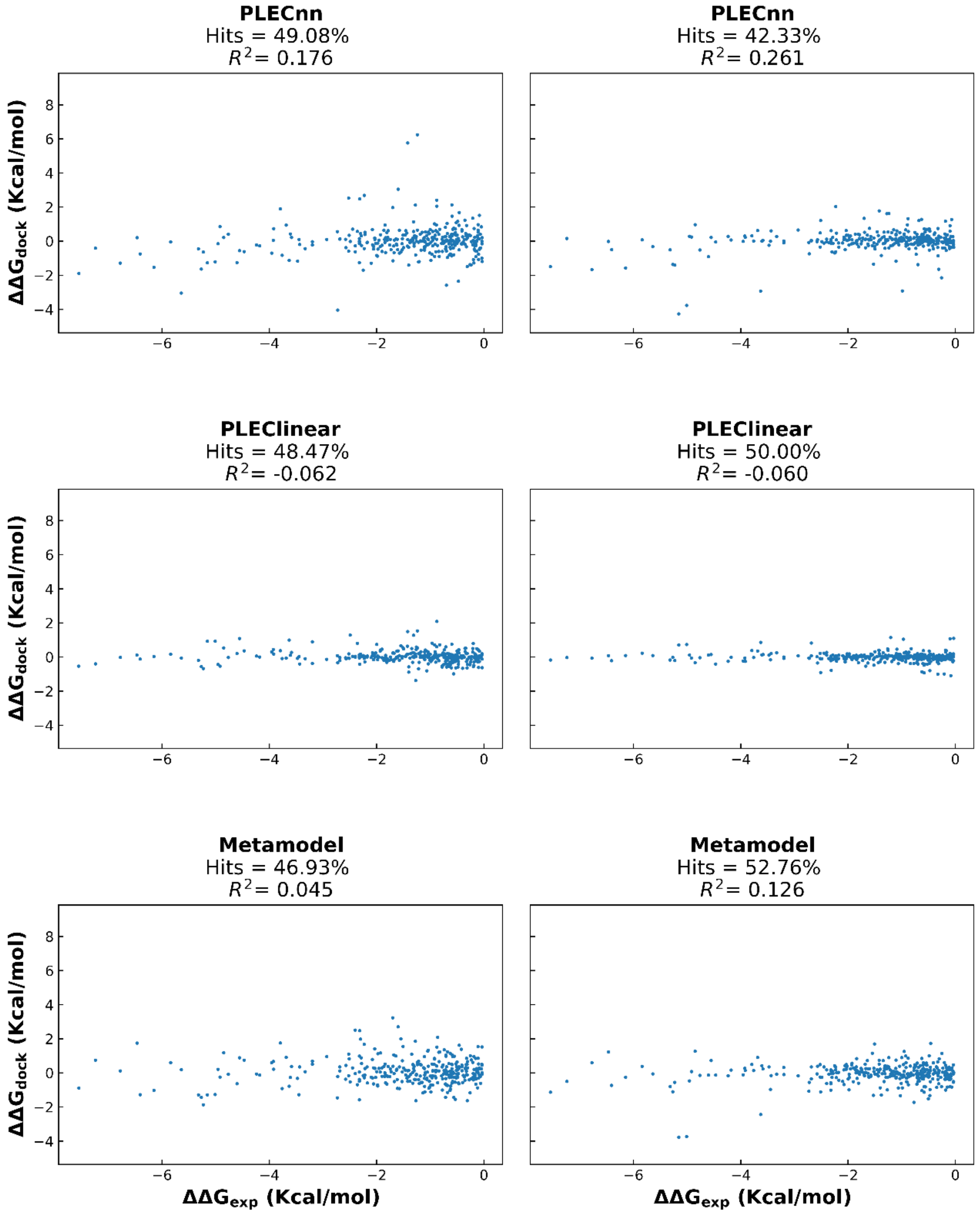


*Figure S9 - Continued. (Page 4 of 4).*
